# Binding of AP endonuclease-1 to G-quadruplex DNA depends on the N-terminal domain, Mg^2+^ and ionic strength

**DOI:** 10.1101/2021.08.25.457676

**Authors:** Aaron M. Fleming, Shereen A. Howpay Manage, Cynthia J. Burrows

**Affiliations:** Department of Chemistry, University of Utah, 315 S. 1400 E., Salt Lake City, UT 84112-0850, United States

**Keywords:** G-Quadruplex, APE1, gene regulation, binding assays, DNA remodeling, pyridostatin

## Abstract

The base excision repair enzyme apurinic/apyrimidinic endonuclease-1 (APE1) is also engaged in transcriptional regulation. APE1 can function in both pathways when the protein binds to a promoter G-quadruplex (G4) bearing an abasic site (modeled with tetrahydrofuran, F) that leads to enzymatic stalling on the non-canonical fold to recruit activating transcription factors. Biochemical and biophysical studies to address APE1’s binding and catalytic activity with the vascular endothelial growth factor (*VEGF*) promoter G4 are lacking, and the present work provides insight on this topic. Herein, the native APE1 was used for cleavage assays, and the catalytically inactive mutant D210A was used for binding assays with double-stranded DNA (dsDNA) versus the native G4 or the G4 with F at various positions, revealing dependencies of the interaction on the cation concentrations K^+^ and Mg^2+^ and the N-terminal domain of the protein. Assays in 0, 1, or 10 mM Mg^2+^ found dsDNA and G4 substrates required the cation for both binding and catalysis, in which G4 binding increased with [Mg^2+^]. Studies with 50 versus physiological 140 mM K^+^ ions present showed that F-containing dsDNA was bound and cleaved by APE1; whereas, the G4s with F were poorly cleaved in low salt and not cleaved at all at higher salt while the binding remained robust. Using Δ33 or Δ61 N-terminal truncated APE1 proteins, the binding and cleavage of dsDNA with F was minimally impacted; in contrast, the G4s required the N-terminus for binding and catalysis. With this knowledge, we found APE1 could remodel the F-containing *VEGF* promoter dsDNA→G4 folds in solution. Lastly, the addition of the G4 ligand pyridostatin inhibited APE1 binding and cleavage of F-containing G4s but not dsDNA. The biological and medicinal chemistry implications of the results are discussed.

**TOC graphic:** 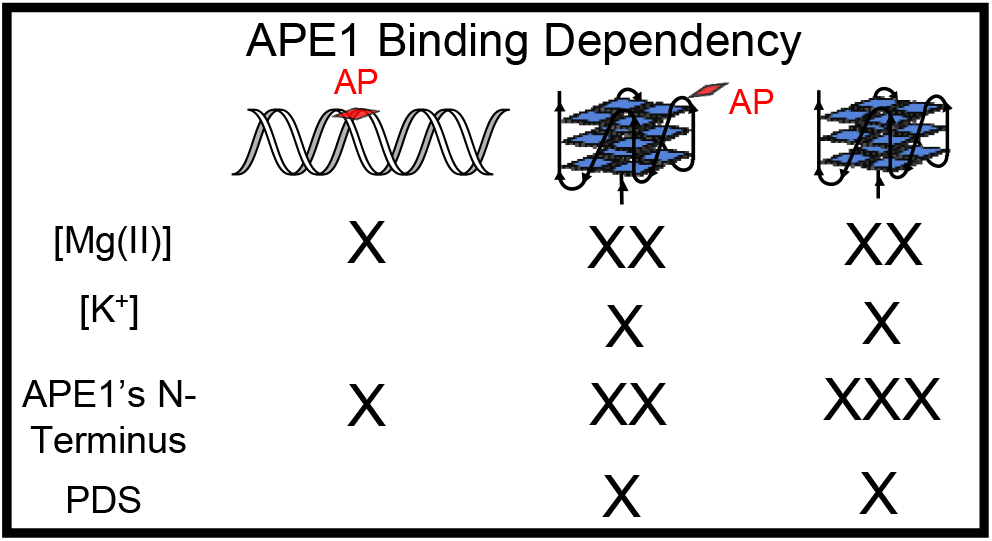

## Introduction

Interactions between DNA and proteins are essential for driving the faithful execution of genomic processes such as the repair of damaged DNA nucleotides and transcriptional regulation. Oxidative stress is a cellular situation in which damage to DNA increases, and transcription of many genes is modulated in response to cellular damage.^1,2^ Contemporary studies have shown diversity in the functions of DNA repair proteins in that some are also actively engaged in transcriptional regulation;^3-9^ these include 8-oxoguanine DNA glycosylase I (OGG1) and apurinic/apyrimidinic endonuclease-1 (APE1).^10,11^ However, the DNA-protein interactions regarding this crossover from repair to regulation are poorly understood, particularly for APE1.

The role of APE1 in transcriptional regulation has been proposed to operate by a few different pathways. Beyond APE1 functioning as an endonuclease in DNA repair, it is a redox effector factor (REF-1) found to reduce transcription factors such as AP-1, HIF-1α, or p53 to their reduced and activated states for DNA binding and gene regulation.^12-16^ APE1 is proposed to stimulate transcription factor loading on DNA via transient cooperative binding to induce a conformational change of the helix.^17^ Another proposed pathway involves APE1 binding to gene promoter sequences bearing an abasic site (AP) substrate and stalling because of the non-canonical structure;^11,15^ In this pathway, the nuclease activity is attenuated allowing recruitment of activating transcription factors such as AP-1 or HIF-1α for gene activation. The cellular finding of APE1 and transcription factor binding to DNA at the same loci observed by ChIP analysis supports the co-binding proposal for gene regulation.^15^ This has been best documented for activation of the vascular endothelial growth factor *VEGF* gene, but there remain questions regarding why stalling of APE1 occurs in the promoter to regulate transcription as opposed to the repair process continuing by the multifunctional endonuclease (Figure 1). We have proposed a role for G-quadruplex folding to explain the switch from APE1 acting as an endonuclease to that of a redox effector for transcription factor recruitment.

**Figure 1.**
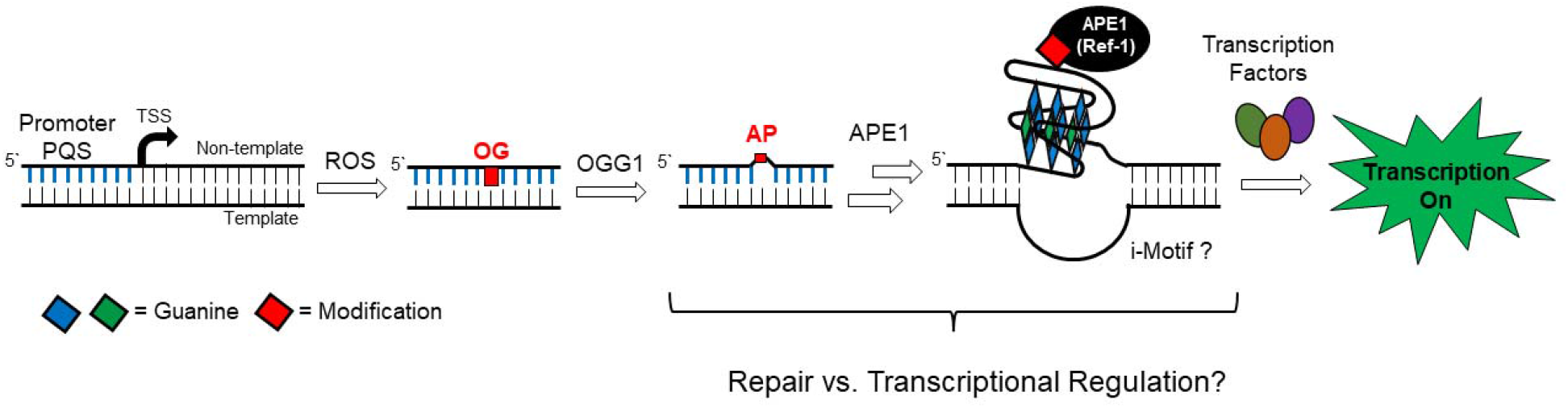
The multifunctional nuclease APE1 is a member of the base excision repair pathway and can function in transcriptional regulation. The present work asks what molecular properties of the DNA and protein lead to the crossover from one function to the other?

Our prior work for the proposal of APE1 binding DNA for gene regulation was derived from chemically-defined plasmid DNA sequences studied in cellulo, and it was based on two features common to G_n_ (n ≥ 3) sequences in DNA.^1,11,18^ First, stacking of two or more adjacent guanines in the DNA duplex lowers the redox potential of G’s followed with a G on their 3’ side, and such sites can be targeted at a distance via long-range charge transport, ultimately forming 8-oxo-7,8-dihydroguanosine (OG) in the GGG track.^19-22^ The OG in double-stranded DNA (dsDNA) then becomes a target for initiation of base excision repair.^23^ Secondly, G-rich sequences such as those found in human gene promoters have a greater propensity to adopt non-canonical structures including G-quadruplex (G4) folds that can function as hubs for transcription factor loading.^24,25^ Thus, remote oxidation of dsDNA can result in OG formation focused in G-rich gene promoters followed by the BER glycosylase OGG1 targeting this site. The AP lesion so formed destabilizes dsDNA which may be the trigger for refolding to a G-quadruplex (G4) (Figure 1).^26,27^

G-quadruplex folds occur in DNA sequences that have at least four closely spaced tracks of three (or more) Gs. Three-layer G4s are stabilized by coordination of two K^+^ ions, the dominant cellular cations, and each G-track is connected by one or more nucleotides that form single-stranded loops arranged in topologies that can orient the 5’-3’ G tracks either parallel or anti-parallel. In many instances in human promoters, there exist extra G nucleotides that can reside in the loops or that are part of a fifth G-run. These extra G nucleotides can aid in maintaining G4 folding even when an essential core G nucleotide is chemically modified, such as being oxidized to an OG lesion.^28^ As a consequence, the AP-bearing G4 is a stable structure. We propose that this G4 structure allows docking of APE1 followed by transcription factor recruitment for gene regulation (Figure 1).^1,2^

There remain many unknowns regarding the biochemistry and biophysics of APE1 interacting with G4s, and specifically promoter G4s, that may determine repair versus regulation by this protein. For instance, APE1 endonuclease activity assays monitoring AP cleavage showed low yields in the human telomere,^29-31^ *c-MYC*^32^ and *NEIL3*^33^ G4 folds in concentrations of K^+^ generally studied below physiological concentrations, and no data have been reported for APE1 activity on an AP-containing *VEGF* G4. The binding constant for APE1 interacting with the native human telomere G4 folds in Na^+^ or K^+^ solutions in which antiparallel and hybrid folds, respectively, have been measured;^30,31^ in contrast, binding constants with predominantly parallel-stranded promoter G4s with and without AP sites have not been evaluated. The APE1 protein requires Mg^2+^ as a critical cofactor for catalysis and product release,^34-36^ but variations in Mg^2+^ concentrations have not been studied with APE1 interacting with G4s. The APE1 protein has an intrinsically disordered N-terminal domain, and its role in binding and influencing the endonuclease activity were reported for the human telomere sequence,^30^ but this is not known for promoter parallel-stranded G4s. Finally, it is not known whether APE1 can remodel duplex DNA to G4-folded DNA to generate the structural hub for the recruitment of transcription factors. The work herein provides new molecular insights for APE1 interacting with promoter G4s, addressing these unknowns. The details of the present study aid in understanding how APE1 functions as both a DNA repair and transcriptional regulatory protein, whereby the crossover from repair to regulation can occur in G-rich gene promoter sequences that can adopt G4 folds.

## Results and Discussion

### 2.1. Characterization of the VEGF G4s studied

The *VEGF* promoter DNA strands were made by solid-phase synthesis with the tetrahydrofuran (F) analog of AP at positions previously found to be prone to oxidative damage (Table 1).^28^ The F analog, unlike an AP, is chemically stable and both are equally good substrates for APE1.^37^ The *VEGF* potential G-quadruplex sequence (PQS) has five G runs in which the four on the 5‵ side adopt the principal wild-type fold in solution.^38^ For the native *VEGF* G4 studies, only the principal fold was studied (*VEGF*-4). When a modification such as an F is present, the fifth G track can swap with the damaged G run to maintain the fold,^28^ and to study this, both four- and five-track *VEGF* sequences were prepared with the F modification. To interrogate the impact of a lesion in a loop position, an F was placed at position 12 (*VEGF*-4 F12 and *VEGF*-5 F12), and to study a core lesion position, an F analog was synthesized at position 14 (*VEGF*-4 F14 and *VEGF*-5 F14). The folding and stabilities of the *VEGF* sequences under the initial endonuclease and binding conditions were first evaluated.

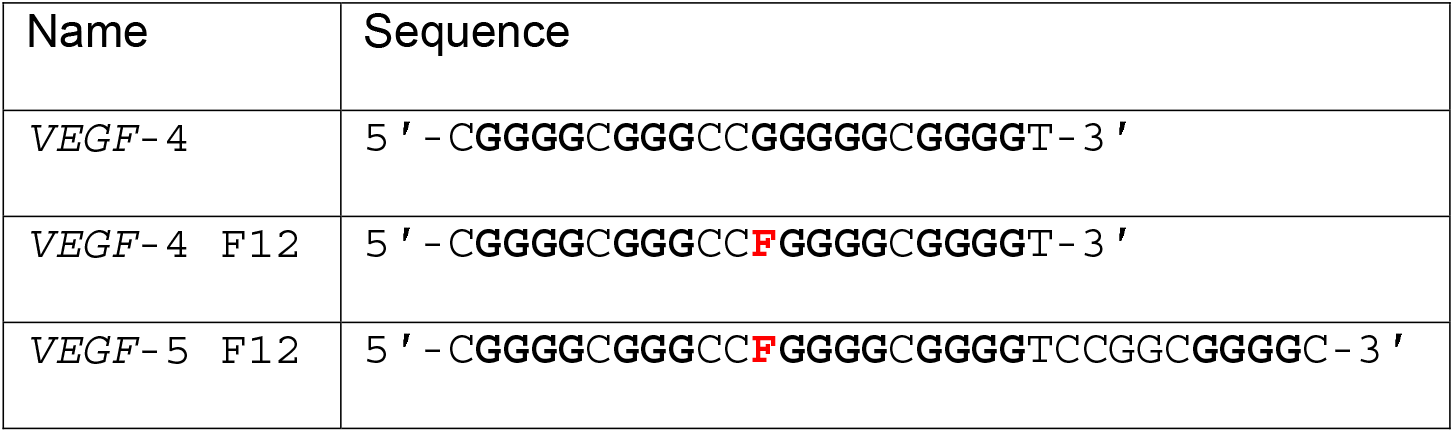
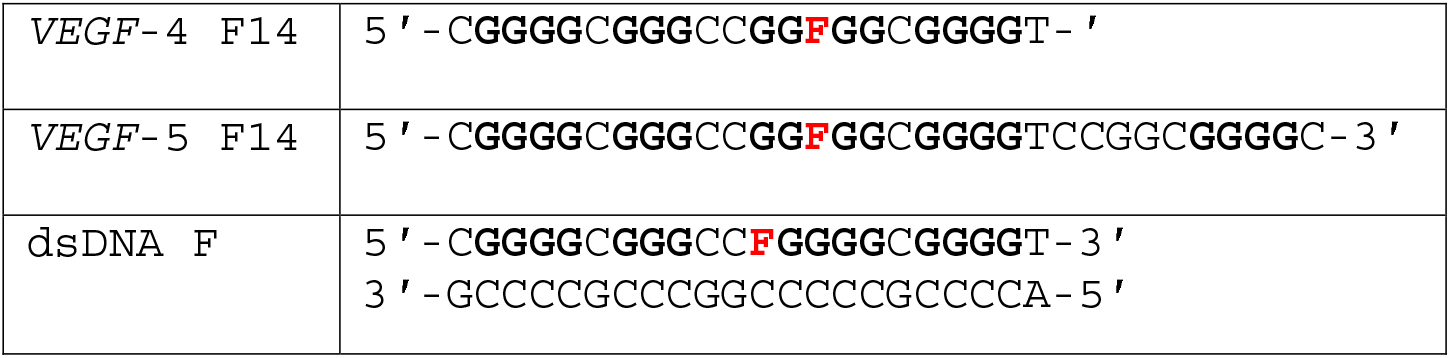

As a consequence of APE1 requiring Mg^2+^ as an essential cofactor for catalysis,^34,35^ the G4 folding studies were conducted with and without Mg^2+^ present to determine whether the G4 topology was altered. The *VEGF* sequences were allowed to fold in APE1 analysis buffer that was selected based on literature reports (20 mM Tris or NaP_i_ buffer pH 7.4 at 22 °C and 50 mM KOAc) with and without 10 mM Mg(OAc)_2_.^32^ The folded G4s were then analyzed by circular dichroism (CD) spectroscopy and thermal melting analysis (*T*_*m*_) to determine the global G4 topologies, their stabilities, and the impact of Mg^2+^ on these two features. Structural control studies were conducted in Li^+^ buffers that do not permit G4 folding.^39^ First, the CD analysis in buffers with Li^+^ ions found the sequences failed to adopt G4 folds based on the low intensity and flat CD spectra obtained (Figures 2A-2E gray lines). These findings suggest the sequence are likely random coils in Li^+^ buffer based on comparisons to the literature.^39^ The native *VEGF*-4 sequence folded in K^+^-ion buffers produced CD spectra with a positive rotation at ∼ 260 nm and negative rotation at ∼240 nm that was not impacted by Mg^2+^ (Figures 2E black vs. red lines). These spectra are good examples of a parallel-stranded G4 fold that is not sensitive to the presence of Mg^2+^.^40^

**Figure 2.**
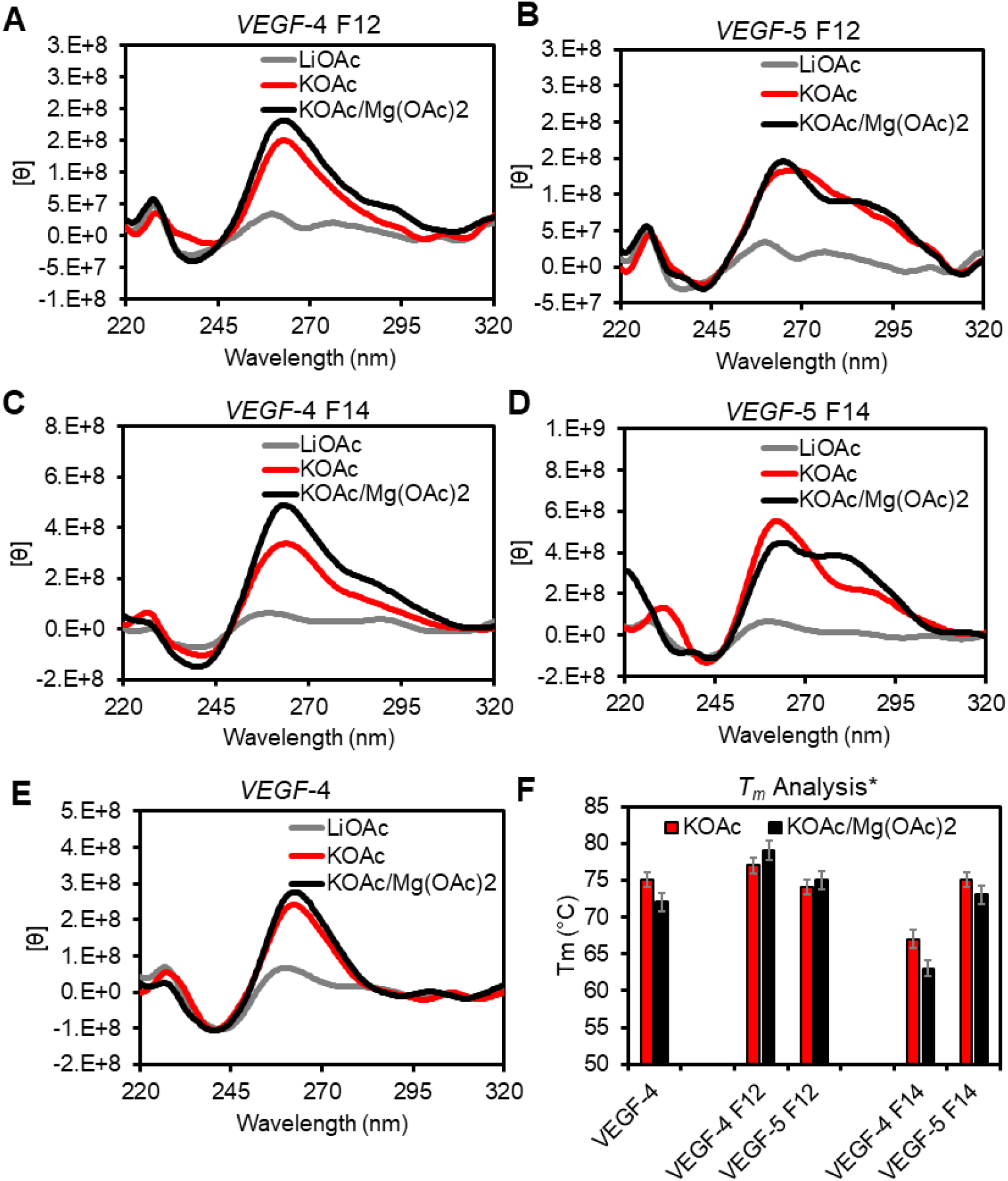
Analysis of the impact of K^+^ and Mg^2+^ cations on the folding and stability of the *VEGF* G4s. (A-E) Inspection of G4 folding by CD spectroscopy in different buffer systems at 22 °C. (F) A plot of the *T*_*m*_ values measured and the respective errors from three independent analyses in the different buffer systems. The buffer and salt compositions represented by the labels are as follows: LiOAc = 20 mM Tris pH 7.4 at 22 °C with 50 mM LiOAc. KOAc = 20 mM Tris pH 7.4 at 22 °C with 50 mM KOAc; KOAc/Mg(OAc)_2_ = 20 mM Tris pH 7.4 at 22 °C 50 mM KOAc and 10 mM Mg(OAc)_2_. *The *T*_*m*_ values were recorded in 20 mM KP_i_ buffer at pH 7.4.

The F-containing G4 folds produced subtle but distinct differences compared to the native *VEGF* G4 studied in the same buffers (Figures 2A-2E). They all had rotation maxima and minima at ∼260 and ∼240 nm, respectively, supporting parallel-stranded G4 folds.^40^ In addition, all spectra recorded had shoulders of positive rotation at wavelengths >280 nm that suggests a subpopulation of antiparallel-folded G4s are also in solution.^40^ When Mg^2+^ ions were added at 10 mM concentration before annealing in K^+^ buffers, no significant changes to the CD spectra were observed for the four-track *VEGF*-4 F12 and *VEGF*-4 F14 sequences, as well as the five-track *VEGF*-5 F12 sequence (Figures 2A, 2B, and 2C black vs. red lines). In contrast, when the fifth G-track was present in the *VEGF*-5 F14 with a core modification that engages the fifth G-track most efficiently, Mg^2+^ produced a significant shoulder at the long-wavelength region with a stronger intensity that suggests a greater amount of antiparallel stranded fold in solution (Figure 2D black vs. red lines). Finally, thermal melting studies used an established protocol to measure the *T*_*m*_ values by monitoring the unfolding process via the decay in UV-vis absorption intensity at 295 nm.^41^ The *T*_*m*_ values for the G4 folds were revealed to be >60 °C and only showed minor differences in the presence of Mg^2+^ (Figure 2F). The most interesting observation regarding the *T*_*m*_ values was the in the G4s with a core lesion at position 14; in the four-track sequence (*VEGF*-4 F14), the *T*_*m*_ value was ∼10 °C lower than the native sequence but when the fifth G track was present, stability was recovered. This is consistent with our prior studies showing engagement of the fifth G-run to stabilize the fold when a lesion resides in a core position.^11,28^

In summary, the native *VEGF*-4 sequence adopts a parallel-stranded G4 under the conditions of the study, a topology consistent with the NMR structural results for this sequence.^38^ The F-containing *VEGF* G4s provided CD spectra that identify both parallel- and antiparallel-folded G4s are present in solution (Figure 2A-E). Extinction coefficients for the CD intensities for each fold type are not known, and therefore, the ratio of parallel to antiparallel cannot be determined. The *T*_*m*_ values measured for all G4 folds were much greater than the 22 or 37 °C temperature at which the binding and activity assays were conducted, respectively, and any differences in the values will not likely impact the results of the experiments that followed.

### 2.2 APE1 endonuclease activity assays for cleavage of the VEGF dsDNA and G4s

Prior reports have demonstrated that the endonuclease activity of APE1 is attenuated when an F is found in the hTelo,^29-31^ *NEIL3*,^33^ or *c-MYC*^*32*^ G4 folds with position dependency on the cleavage yield; however, this analysis has not been reported for the *VEGF* G4 sequences with an F at loop or core positions. The present studies monitored APE1-catalyzed cleavage at an F by separating the reactants from the products on a polyacrylamide gel electrophoresis (PAGE) experiment using ^32^P 5′-labeled DNA strands to visualize and quantify the bands (Figure 3A). The first reactions were conducted using previously established reaction buffers and salts (20 mM Tris pH 7.5 at 37 °C, 50 mM KOAc, 10 mM Mg(OAc)_2_, and 1 mM DTT).^32^ The reactions were conducted under steady-state reaction conditions with 10 nM DNA and 3 nM APE1. First, the dsDNA F with a C nucleotide opposite the lesion was used as the positive control substrate to monitor a 60 min time-course product production profile catalyzed by APE1 (Figure 3B). The yield was >90% after 5 min verifying a robust reaction in this sequence context consistent with prior studies.^36,42,43^

**Figure 3.**
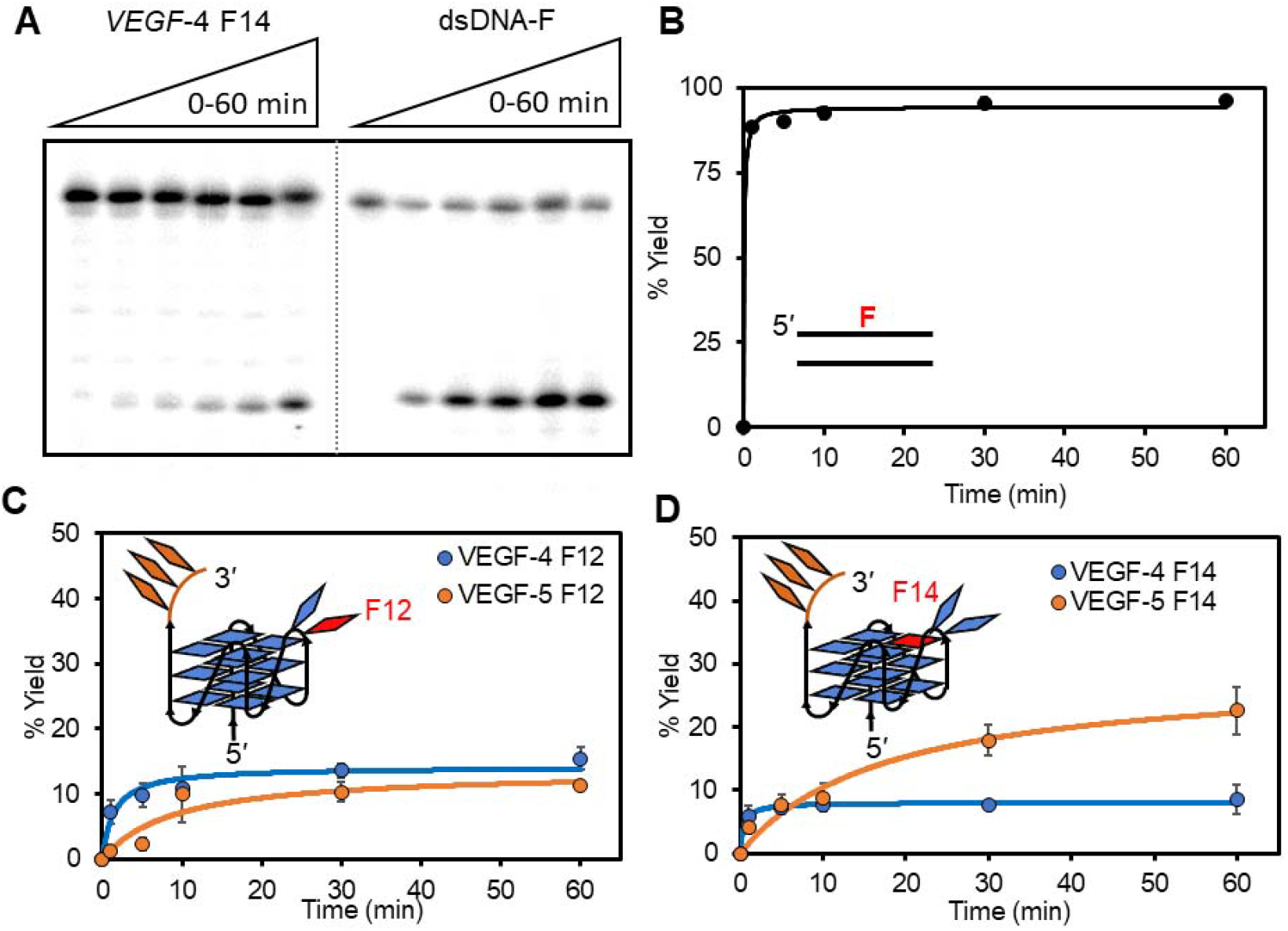
Monitoring the APE1-catalyzed cleavage of an AP site in the *VEGF* sequence by PAGE analysis to identify position- and context-dependent yields. (A) Example time-dependent PAGE analysis of APE1-catalyzed strand scission at an F site in the dsDNA or G4 contexts. The yield vs. time profiles for cleavage in the (B) dsDNA context with F at positions 12 (dsDNA F), as well as four- and five-track G4 contexts with F at position (C) 12 or (D) 14. The reactions were conducted by preincubating 10 nM DNA in 20 mM Tris pH 7.5 (37 °C) with 50 mM KOAc, 10 mM Mg(OAc)_2_, and 1 mM DTT present at 37 °C for 30 min. After the preincubation, APE1 was added to a 3 nM final concentration, and the reactions were allowed to progress to the desired time and then quenched by adding a stop solution and heating at 65 °C for 20 min before PAGE analysis.

The time-course reaction profiles for the F-containing *VEGF* G4s cleaved by APE1 showed that the yields never surpassed ∼20%, and the value measured was highly dependent on the location of the lesion and whether the fifth G track was present or not. For F12 with a G4 loop lesion in the four-track G4 (*VEGF*-4 F12), the APE1-catalyzed cleavage yield was nearly 15% at 10 min and slightly increased after a 60 min reaction; the same lesion in the five-track fold (*VEGF*-5 F12) more slowly progressed to finally achieve a cleavage yield of ∼15% after 60 min (Figure 3C). In contrast, the sequences with a core F at position 14, the four-track fold (*VEGF*-4 F14) was cleaved by APE1 at 10 min to a yield of ∼8% that did not change up to the 60 min cutoff (Figure 3D). Interestingly, when the fifth G-track was present, cleavage of the core F slowly reached a final yield of ∼20% after a 60 min reaction (Figure 3D). These data verify APE1 cleavage of an F site in the *VEGF* G4 context with either four or five G tracks is highly attenuated compared to cleavage in the dsDNA context. The final yield for APE1-catalyzed cleavage of an F in the *VEGF* G4 context is dependent on the location of the lesion, similar to findings for F cleavage by this enzyme in the hTelo and *NEIL3* G4 folds.^29,30,33^ In these studies, the fifth G track slowed the rate of the endonuclease reaction that likely results from the increased dynamics of the G4 fold when the extra domain is present.

The enzyme assays described were conducted under steady-state reaction conditions ([DNA] > [APE1]) that could result in product inhibition for the G4 reaction contexts leading to low overall yields compared to the dsDNA context; this has been observed for APE1 in other reactions.^36^ To probe the possibility of product inhibition for the reactions, experiments were then conducted under single-turnover conditions ([DNA] = [APE1] = 10 nM) to observe that the 60-min yield remained the same (Figure S1). Many other variations in the reaction setup were attempted to find the yield still did not change (Figure S1). This led us to question whether APE1 was even capable of binding the F-containing, and predominantly parallel-stranded, *VEGF* G4 substrates that led to the poor endonuclease yields (Figures 2 and 3).

### 2.3 Cation dependency for binding of APE1 to the F-containing VEGF dsDNA and G4s

The full-length native APE1 was used for the initial studies. Because Mg^2+^ is a key cofactor required for the endonuclease activity of APE1 and cleavage would interfere with binding, the initial studies were conducted in the presence of excess EDTA and no added Mg^2+^. Control activity assays found under the saturating EDTA conditions employed, no APE1 cleavage was observed (Figure 3A; displayed as no reaction (n.r.) for what would be orange bars). Typical methods to monitor and quantify DNA-protein binding interactions include electrophoretic mobility shift assays (EMSAs) or surface plasmon resonance (SPR). Attempts to study the APE1-G4 binding interaction by EMSAs produced bands in the wells that were not suitable for quantification; in the SPR analysis of the interaction, the nonspecific interaction of APE1 with the sensor chip was too high to yield reliable data for analysis (Figure S2). Therefore, fluorescence anisotropy was applied to monitor and quantify the APE1-G4 binding interaction, an approach previously applied to quantify protein-G4 binding constants and APE1-dsDNA binding constants.^44,45^ There exist a few drawbacks to this approach: (1) Either the DNA or protein must be labeled with a fluorophore; FAM-labeled DNA was used in the present work. CD studies verified that FAM labeling of the sequences did not influence the G4 folding. (Figure S3). (2) Another challenge with fluorescence anisotropy is that binding is determined by a change in fluorophore emission from the DNA as the titrated protein binds the DNA leading to a plateau in the emission; thus, the data do not report on the absolute quantity of bound DNA but instead give a profile of those for which binding occurs. With this known limitation, binding between APE1 and the *VEGF* G4s was studied by fluorescence anisotropy.

The fluorescence anisotropy values (r) obtained from a titration of APE1 from 0-5000 nM into a solution of 100 nM DNA were plotted against the log[APE1] to produce sigmoidal curves that were fit to the appropriate Hill equation; fitting of the data allowed determination of the dissociation-binding constant (*K*_*D*_) and the Hill coefficients reporting the molecularity of the interaction (Figure 4B).^46^ Without Mg^2+^ present, native APE1 bound the dsDNA F with a *K*_*D*_ value of 50 ± 4 nM (Figure 4C orange). This value is slightly higher than reported with a 42-amino acid N-terminal truncated form of APE1 binding an F-containing dsDNA,^44^ which is the likely source of the difference. In contrast, binding to dsDNA without an F site gave a *K*_*D*_ > 5000 nM that is the maximum WT APE1 concentration studied (Figure 4B gray line). These values on dsDNA substrates verify APE1 has robust binding to dsDNA-F and negligible binding to DNA without the abasic substrate.

**Figure 4.**
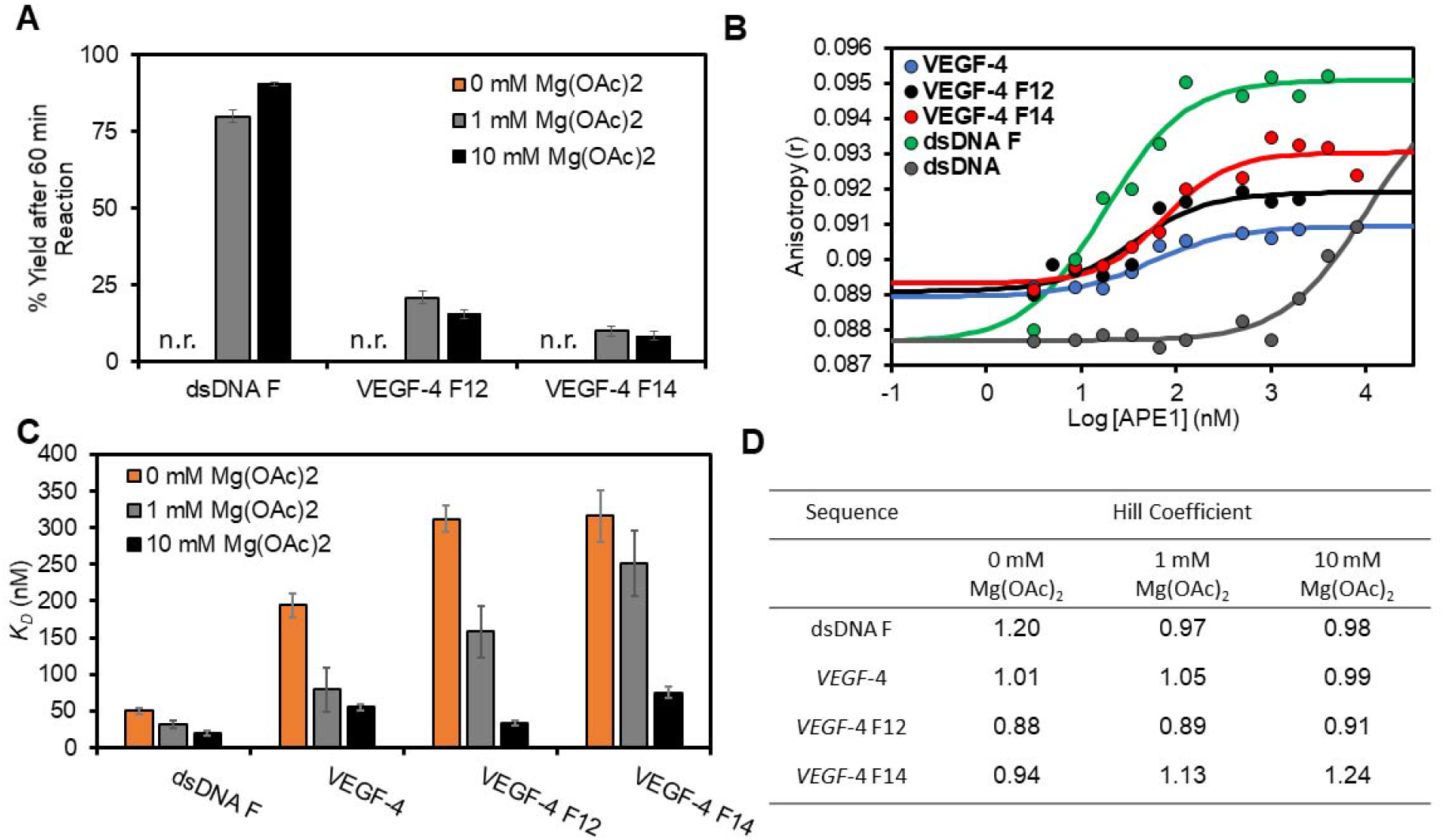
Endonuclease activity and fluorescence anisotropy binding assays for the interaction between APE1 and the *VEGF* PQS dsDNA or G4 folds demonstrate a Mg^2+^ dependency. (A) The [Mg^2+^]-dependent activity for native APE1 on the F-containing *VEGF* substrates was measured after 60 min. (B) Binding plots with log[APE1] vs. anisotropy (r) for the analysis without Mg^2+^ present. (C) The *K*_*D*_ values were obtained from fitting the sigmoidal binding curves with the appropriate Hill equation. The 0 mM Mg^2+^ data were obtained from WT APE1, while the 1 and 10 mM Mg^2+^ data were obtained using the catalytically inactive D210A-APE1 mutant. The errors represent the standard deviation of triplicate trials. (D) Table of Hill coefficients obtained from the fits. The studies were conducted in 20 mM Tris, 50 mM KOAc and 0, 1, or 10 mM Mg(OAc)_2_, 1 mM DTT at pH 7.5 for the activity assays at 37 °C and pH 7.4 for the binding assays at 22 °C. n.r. = no reaction

Binding assays with G4 DNAs produced *K*_*D*_ values that are larger than the one observed with dsDNA F. The *VEGF*-4 F12 containing a loop lesion produced a 310 ± 30 nM *K*_*D*_ value (Figure 4C orange), and *VEGF-*4 F14 G4 with a lesion in a core position had a 320 ± 80 nM *K*_*D*_ value for the interaction (Figure 4C orange). Lastly, binding of native APE1 to the *VEGF*-4 G4 without an F displayed a *K*_*D*_ value of 190 ± 50 nM for the binary complex (Figured 4C orange). Finally, the Hill coefficients found by the fits were all around 1 (0.88-1.24) suggesting a one-to-one interaction between WT APE1 and the *VEGF* sequences under the conditions of the present experiments (Figure 4D). These studies without Mg^2+^ present suggest the binding between APE1 and the G4s have a sixfold greater *K*_*D*_ value for the quadruplex compared to the dsDNA F substrate.

Beyond being an essential cofactor, Mg^2+^ may also have an effect on substrate binding;^34^ however, to study this divalent metal ion in a binding assay, the catalytic function must be abolished. The proposed mechanism for APE1 to cleave the phosphodiester 5‵ to an AP uses Asp210 to activate the water molecule that attacks the phosphodiester;^47,48^ and therefore, mutation of Asp210 to Ala210 renders the protein catalytically inactive even when Mg^2+^ is present;^49^ additionally, Asp210 is not involved in Mg^2+^ coordination by the protein. The D210A mutant of APE1 (D210A-APE1) was then produced by standard recombinant methods and used to probe the binding between APE1 and the *VEGF* substrates in the standard buffer with 10 mM Mg^2+^ present. Before commencing the binding studies, we verified that the mutant protein could not cleave an F-containing dsDNA with 10 mM Mg^2+^ present (Figure S4). The D210A-APE1 interaction with dsDNA F showed a *K*_*D*_ value of 20 ± 4 nM, which is 2.5-fold lower than without Mg^2+^ present (Figure 4C black); further, the lower value is similar to that of prior studies regarding APE1 binding to F-containing dsDNA.^35,44,50,51^ The binary complex between D210A-APE1 and the unmodified *VEGF*-4 G4 had a *K*_*D*_ value of 55 ± 4 nM that is 3.5-fold lower than studies without Mg^2+^ (Figure 4C black). The *VEGF*-4 F12 sequence had *K*_*D*_ values of 34 ± 3 nM when Mg^2+^ was present that corresponds to a reduction by 9.2-fold relative to the experiments without Mg^2+^ (Figure 4C black). The D210A-APE1 interaction with *VEGF*-4 F14 in the presence of Mg^2+^ dropped the *K*_*D*_ value to 75 ± 8 nM that is 4.2-fold lower than found without Mg^2+^ (Figure 4C black). The binding constants for the D210A-APE1 to the *VEGF* G4 substrates in Mg^2+^ were significantly lower than the interaction between WT APE1 and the same DNAs in the absence of Mg^2+^. Moreover, in the presence of Mg^2+^, the APE1 binding to F-containing dsDNA and G4 folds were similar, suggesting that APE1 could favorably bind the non-canonical G4 folds in cells.

Studies regarding APE1-G4 activity and binding are generally conducted in lower ionic strength solutions with K^+^ concentrations at ∼ 50 mM and Mg^2+^ at ∼ 10 mM.^32^ The intracellular concentration of K^+^ is ∼140 mM, the highest value for a monovalent cation, and it is the cellular cation that stabilizes G4 folds to the greatest extent;^52^ an interesting contrast is that Li^+^ ions are not compatible with G4 folding,^52^ which was reiterated in the present work (Figure 2A-E gray lines) and will be used below to address the G4 fold vs. G-richness of the strands for interacting with APE1. The intracellular concentration of Mg^2+^ can reach 10-20 mM depending on the cell type, and the free Mg^2+^ concentration is typically ∼1 mM.^53,54^ This knowledge sets the stage for a set of cation-dependent experiments regarding the binding and activity of APE1 with the *VEGF* sequences.

In the first studies, the Mg^2+^ concentration was dropped tenfold to 1 mM while maintaining all other factors constant to determine the [Mg^2+^]-dependency in APE1 activity and its binding constants with the *VEGF* substrates. The APE1 cleavage activity on the dsDNA F substrate at 1 mM gave a ∼80% yield, a value slightly less than when 10 mM Mg^2+^ was present (Figure 4A gray vs. black bars). This is consistent with prior work that found the activity of APE1 on dsDNA is Mg^2+^ dependent and close to maximal activity at ∼10 mM.^55^ In contrast, the G4 substrates *VEGF*-4 F12 and *VEGF*-4 F14 were cleaved by APE1 in yields at 1 mM Mg^2+^ similar to those in 10 mM Mg^2+^ (Figure 4A gray); this observation demonstrates there is not a strong Mg^2+^ dependency for APE1 cleavage of an F in the *VEGF* G4 context. As for APE1 binding the *VEGF* dsDNA F, *VEGF*-4, *VEGF*-4 F12, or *VEGF*-4 F14, the *K*_*D*_ values with 1 mM Mg^2+^ present were between those for 0 and 10 mM Mg^2+^ values previously discussed (Figure 4C gray). These intermediate values clarify the trend that APE1 binding is Mg^2+^ dependent and is achieved with greater affinity as reflected by a lower *K*_*D*_ value as the divalent metal ion concentration increases. Lastly, to reiterate, the Mg^2+^ effect on binding was most significant for G4 folds. On the other hand, the activity of APE1 to cleave an F in the *VEGF* G4s is less Mg^2+^ dependent than the binding.

The next studies evaluated the impact on APE1 interacting with the *VEGF* sequences when the monovalent cation K^+^ was at physiological intracellular concentrations (140 mM). The activity assays revealed that at higher K^+^ concentrations the APE1 cleavage yield for dsDNA F was 75% while cleavage was not detectable for the F-containing *VEGF*-4 sequences (Figure 5A green). These cleavage yields are dramatically lower than those measured when the [K^+^] = 50 mM (Figure 5A black). To test whether the lower activity results from compromised binding, the *K*_*D*_ values were then determined when the [K^+^] = 140 mM. The *K*_*D*_ value measured for the D210A-APE1-dsDNA F interaction at 140 mM K^+^ ion was slightly higher than that found at the 50 mM concentration (27 vs. 20 nM, respectively; Figure 5B green vs. black). In contrast, the unmodified *VEGF*-4 substrate was bound by D210A-APE1 with a lower *K*_*D*_ value at the higher concentration of K^+^ ions than the lower value studied (20 vs. 55 nM; Figure 5B green). Similarly, the *VEGF*-4 F14 G4 displayed a lower *K*_*D*_ value in the 140 mM K^+^ sample compared to the 50 mM K^+^ sample (50 vs. 75 nM; Figure 5B green vs. black). Lastly, the *VEGF*-4 F12 G4 had a similar binding constant between the two different concentrations of K^+^ (∼30 nM; Figure 5B). At the higher concentration of K^+^ ions, the binding of APE1 to F in the dsDNA versus G4 contexts were similar; in contrast, the endonuclease activity was somewhat slowed for the dsDNA context and completely abolished in the G4 contexts at the higher ionic strength. These K^+^ concentration-dependent findings may result from the >15 °C *T*_*m*_ values for the G4s previously measured in 140 mM K^+^ buffers.^11,28^ The increased *T*_*m*_ values may decrease the DNA flexibility and prevent APE1 from bending the strand to the 35° angle needed for catalysis.^47^ On the other hand, a less flexible G4 may be a better binding partner with APE1 as described below. Going back to the gene regulation proposal (Figure 1),^11^ these data support the concept that APE1 can bind the promoter F-containing G4 folds tightly without cleaving the strand, resulting in stalling of the enzyme on DNA. More importantly, the binding of APE1 without cleavage at the G4 hub can provide the opportunity to recruit activating transcription factors for gene induction, as previously proposed (Figure 1).^11^

**Figure 5.**
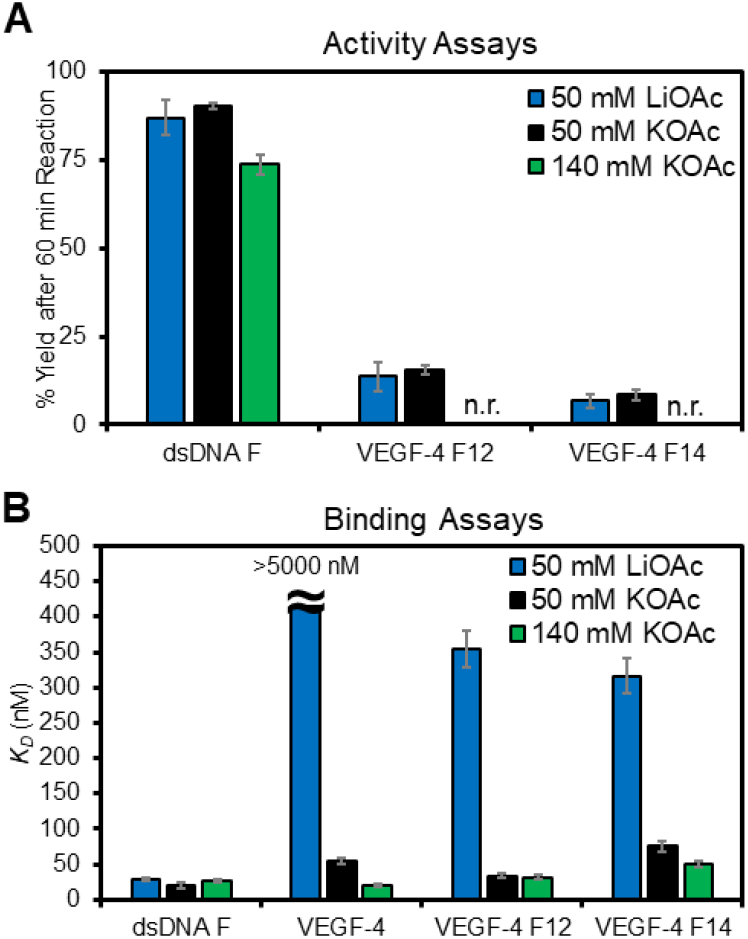
Monovalent cation dependency on the APE1 activity and binding of D210A-APE1 with the *VEGF* PQSs as duplexes or G4s. (A) Endonuclease yields for APE1 cleaving an F in the dsDNA and G4 contexts in 50 mM Li^+^ (blue), 50 mM K^+^ (black), or 140 mM K^+^ buffers (green). (B) Plots of the *K*_*D*_ values measured for D210A-APE1 binding the *VEGF* dsDNA and G4 structural contexts studied. n.r. = no reaction

The final cation-dependent study conducted was in a buffered solution containing Li^+^ ions that are not compatible with G4 folding;^52^ therefore, these data provide a control study for the importance of the G4 fold in the APE1 binding interaction. The G-rich *VEGF* sequences, when annealed in 50 mM Li^+^ buffers, produced CD spectra of low overall intensity without any major defining features compared to those recorded in buffers with K^+^, indicating no G4 folding under these conditions (Figure 2A-E). The APE1 activity in 50 mM Li^+^-containing buffers for the dsDNA F substrate was similar to the yield found in the K^+^-containing buffers at the same ionic strength (Figure 5A blue). A similar result was found for the *VEGF*-4 F12 and *VEGF*-4 F14 G4 substrates in which the yields in Li^+^-containing buffers were very similar to the K^+^-containing buffers of the same ionic strength (Figure 5A blue). This suggests the cleavage yields for G4 folds are similar to that of a single-stranded context.

Next, the binding between D210A-APE1 and the *VEGF* sequences was measured in the Li^+^-containing buffers. In studies with D210A-APE1 binding dsDNA F, the cation identity did not change the *K*_*D*_ value measured (Figure 5B blue). For the unmodified G4 fold *VEGF*-4, no detectable binding was observed in the Li^+^-containing buffers that suggest the protein fails to bind ssDNA without an F substrate (*K*_*D*_ > 5000 nM Figure 5B blue). For the *VEGF*-4 F12 and *VEGF*-4 F14 sequences containing an APE1 substrate F, the *K*_*D*_ values in Li^+^-containing buffers were 360 ± 30 and 320 ± 20 nM, respectively, which are 10.7-fold and 4.2-fold larger than those measured in K^+^-containing buffers (Figure 5B blue vs. black). These cation-dependent studies demonstrate D210A-APE1 binding is significantly enhanced for the *VEGF* G4 folds when an abasic site is present. This further supports the hypothesis that the G4 fold is a structure that is recognized by the APE1 protein to facilitate tight binding.

### 2.4 The fifth G track in VEGF can influence APE1 binding F-containing G4s

We next asked if the presence of a fifth G track that can swap with lesion-containing G runs to maintain the fold would impact APE1 binding to DNAF (Figure 6A).^28,29,33^ The initial studies were conducted with the D210A-APE1 with 10 mM Mg^2+^ present in 50 mM K^+^-containing buffers. The *K*_*D*_ values measured in four-vs. five-track *VEGF* G4s when F was in a loop position, i.e., *VEGF*-4 F12 and *VEGF*-5 F12, produced similar binding constants (∼35 nM; Figure 6B). In contrast, with the core position F that requires the engagement of the fifth track to stabilize the fold (based on *T*_*m*_ analysis (Figure 2F)), the extra G run had a large impact on the binding constant measured. The *VEGF*-5 F14 sequence was bound by D210A-APE1 with a *K*_*D*_ = 36 ± 5 nM while the *VEGF*-4 F14 sequence had a *K*_*D*_ = 75 ± 8 nM, a value that is twofold lower (Figure 6B). These results provide quantitative support for the fifth G-track or “spare tire”^28^ to facilitate APE1 interaction with stable G4 folds bearing an F site. Many of the cation-dependent studies were repeated with the *VEGF*-5 sequences containing F that provided similar conclusions already made with the four-track *VEGF* sequences described in the text (Figure S5). The key finding is the fifth domain when present in the *VEGF* G4 can slightly increase cleavage at an AP site to ∼15-20% yield after 60 min in 50 mM ionic strength buffer (Figure 3C-D), the cleavage yield is not detectable at physiological ionic strength (Figure S5), and APE1 binding can be greatly enhanced with the spare tire domain when the lesion is initially in a core position of the G4 (Figure 6B).

**Figure 6.**
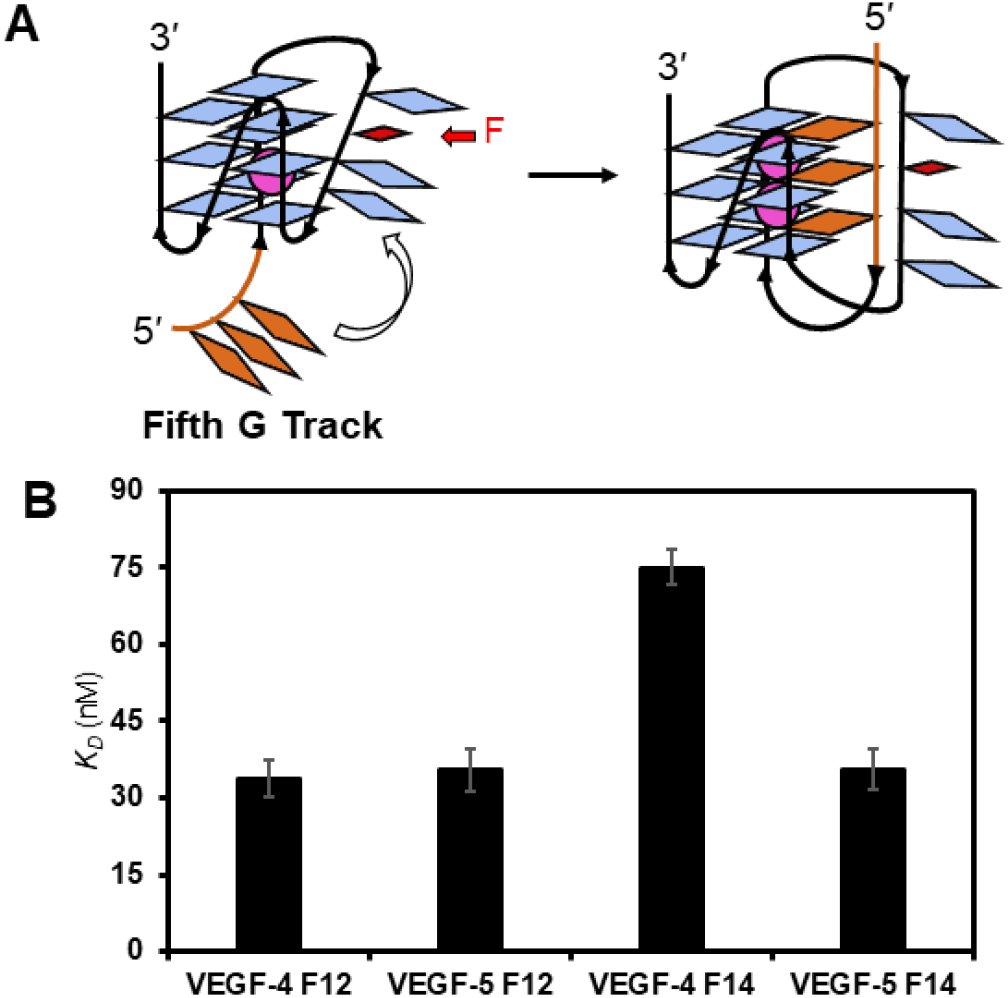
The fifth G track in the *VEGF* G4 can facilitate (A) folding and (B) APE1 binding to F-containing folds.

### 2.5 Analysis of the N-terminal domain of APE1 for binding the VEGF dsDNA and G4s

Beyond Mg^2+^, what are the other essential factors for tight G4 binding by APE1? The APE1 protein has an N-terminal disordered domain of 61 amino acids,^43,56^ in which the first 33 amino acids include many lysine residues. Moreover, removal of the first 33 amino acids from the N-terminus of APE1 was previously found to decrease the activity of F removal from the human telomere G4, as well as binding of APE1 to the Na^+^-folded human telomere sequence.^30^ Therefore, binding and activity assays were conducted with N-terminal truncated APE1 proteins processing the *VEGF* G4s. Studies were used with partial (Δ33) and complete (Δ61) removal of the N-terminal domain of APE1. The activity assays maintained the wild-type active sites of APE1, while the binding studies were conducted with N-terminal truncated D210A-APE1 double mutants produced by standard methods.

Studies to test the dependency of the N-terminal domain for activity on the *VEGF* DNA substrates were conducted in buffered solutions with 50 mM K^+^ ions and 10 mM Mg^2+^ ions present. The APE1 activity assays showed that the dsDNA F substrate yields were the same for APE1 and Δ33-APE1, and decreased by ∼15% for the Δ61-APE1 (Figure 7A). Consistent with prior studies, the absolute APE1 cleavage yields are not altered in dsDNA contexts; however, the prior study found the rate of the reaction is impacted by the loss of the N-terminus.^43^ Similarly, APE1 cleavage efficiency for the four-track *VEGF* sequences *VEGF*-4 F12 and *VEGF*-4 F14 APE1 and Δ33-APE1 were similar in yield (Figure 7A); however, Δ61-APE1 with complete removal of the N-terminal domain was incapable of cleaving the F substrate in the *VEGF* G4 contexts (Figure 7A); even addition of the fifth-domain to the G4 fold failed to be cleaved by the endonuclease (Figure S5). These data suggest the complete N-terminal domain is required to enable endonuclease activity to occur on G4 substrates.

**Figure 7.**
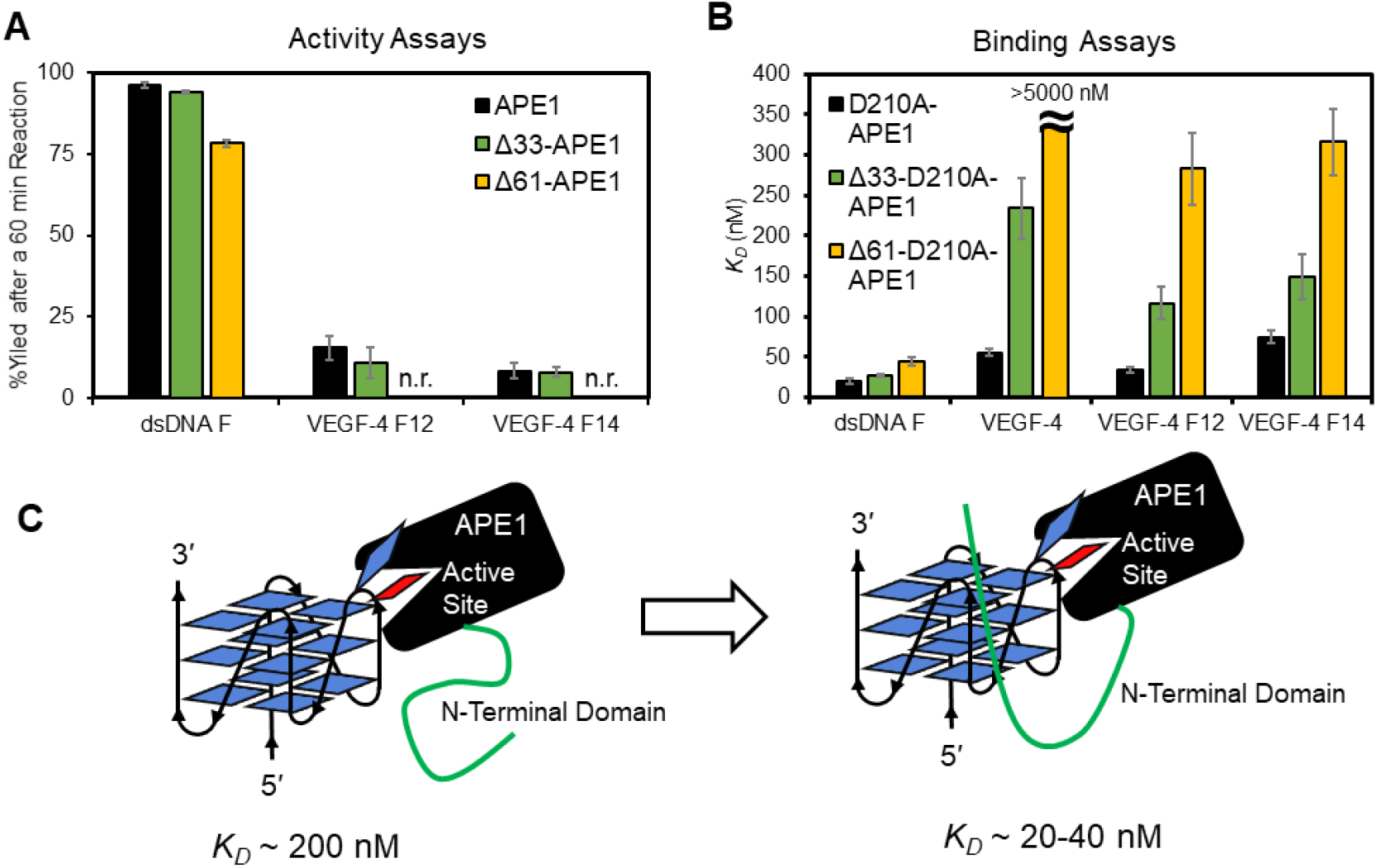
The N-terminal disordered domain of APE1 is required for maximal activity and the lowest *K*_*D*_ values for the protein interacting with the *VEGF* G4s. (A) The cleavage yields after 60 min when full-length APE1 and its N-terminal truncated forms Δ33-APE1 or Δ61-APE1 were allowed to react with an F in dsDNA or the *VEGF* G4 folds. (B) The binding of N-terminal domain truncated and catalytically inactive double mutants (D210A-APE1, Δ33-D210A-APE1, or Δ61-D210A-APE1) with F-containing G4 and dsDNA substrates. (C) Model illustrating the N-terminal domain enables stronger interactions between the APE1 protein and G4 DNA. All analyses were conducted in 20 mM Tris, 50 mM KOAc, 10 mM Mg(OAc)_2_, and 1 mM DTT buffers at pH 7.5 and 37 °C for the activity assays and pH 7.4 and 22 °C for the binding assays. n.r. = no reaction

Binding assays with the N-terminal domain truncated and D210A-APE1 double mutants with the DNA substrates were studied next. The *K*_*D*_ values measured for D210A-APE1 binding to the dsDNA F substrate were slightly impacted by the loss of the N-terminal domain (D210A-APE1: *K*_*D*_ = 20 ± 4 nM; Δ33-D210A-APE1: *K*_*D*_ = 27 ± 3 nM; and Δ61-D210A-APE1: *K*_*D*_ = 44 ± 5 nM; Figure 7B). Complete removal of the N-terminal domain from APE1 increased the *K*_*D*_ value by twofold for dsDNA, a change similar to previous reports for APE1 binding to dsDNA with an F. The wild-type *VEGF* G4 binding was highly impacted by the loss of the N-terminal domain; the D210A-APE1 had a *K*_*D*_ = 55 ± 4 nM, Δ33-D210A-APE1 giving a *K*_*D*_ = 230 ± 40 nM, and the Δ61-D210A-APE1 failed to yield a measurable binding constant (*K*_*D*_ > 5000 nM) in the range studied (Figure 7B). The *VEGF*-4 F12 sequence was bound with lower affinity as the N-terminal tail was successively truncated; the D210-APE1 had a *K*_*D*_ = 34 ± 3 nM, Δ33-D210A-APE1 had a *K*_*D*_ = 120 ± 20 nM, and Δ61-D210A-APE1 had a *K*_*D*_ = 280 ± 50 nM when binding a loop F at position 12 in the G4 (Figure 7B). A similar change was observed for the *VEGF*-4 F14 sequence with a core F in which D210A-APE1 had a *K*_*D*_ = 75 ± 8 nM, Δ33-D210A-APE1 had a *K*_*D*_ = 150 ± 30 nM, and Δ61-D210A-APE1 had a *K*_*D*_ = 320 ± 40 nM (Figure 7B). The trends in the dependency of the N-terminal domain of APE1 binding the five-track *VEGF* G4s *VEGF*-5 F12 and *VEGF*-5 F14 were similar to those observed for the four-track G4s (Figure S6).

Conclusions regarding the impact of the N-terminal domain of APE1 in =binding and cleaving the *VEGF* sequence are many. (1) The N-terminal domain of APE1 is necessary for cleavage of an F in the G4 context in the 50 mM ionic strength conditions, and in contrast, dsDNA F cleavage was not as sensitive to loss of the disordered N-terminal region. (2) The binding of APE1 to F-containing G4 folds is the tightest when the N-terminal domain is present; the active site binding to the F in the G4 context provides a *K*_*D*_ of ∼200 nM based on the Δ61-D210A-APE1 results (Figure 7B), a value that is similar to D210A-APE1 binding the unfolded F-containing G4s in Li^+^ ion conditions (∼300 nM; Figure 5B). Thus, the addition of the N-terminal tails drops the *K*_*D*_ value by 5-to 10-fold depending on the position of the F in the *VEGF* G4 fold (Figure 7C). (3) The binding of APE1 to the native *VEGF* G4 fold is likely entirely through the N-terminal domain, which requires all 61 amino acids because once this domain is removed, no binding was observed (Figure 7B).

### 2.6 Impact of APE1 binding to the VEGF G4s in the presence of a G4 ligand

An approach to probe protein-G4 interactions is through the use of G4 ligands, and this was recently used to identify G4-binding proteins in cells.^57,58^ A common G4 ligand used for such studies is pyridostatin (PDS; Figure 8A), which shows high selectivity for G4 folds over other DNA or RNA structures.^59^ The same approach outlined herein was followed to study the impact of the PDS ligand added at 5 μM for 30 min before following APE1 cleavage activity or binding as described above in buffered solutions with 50 mM K^+^ ions and 10 mM Mg^2+^ ions. The G4 ligand had a minimal impact on the cleavage of the dsDNA F substrate, while in the *VEGF*-4 F12 the yield decreased by more than half, and for *VEGF*-4 F14, no cleavage was detectable (Figure 8B). In the binding assays with D210A-APE1, the interaction with dsDNA F was not significantly perturbed, but the native G4 fold *VEGF*-4 was bound by APE1 with a 12.7-fold greater *K*_*D*_ value (700 nM vs. 55 nM; Figure 8C). The affinity of APE1 for the *VEGF*-4 F12 and *VEGF*-4 F14 G4s decreased by 10.5-fold and 3.7-fold, respectively, when the G4 ligand PDS was present (Figure 8C). The PDS ligand had a similar negative impact on APE1 binding the five-G track *VEGF*-5 F12 and *VEGF*-5 F14 G4s (Figures S6). These data demonstrate that G4 ligands block APE1 binding to parallel-stranded G4s; additionally, this may explain why, in the intracellular studies to identify G4-binding proteins, APE1 was not found in one study and not a major G4 binding partner in the second study.^57,58^

**Figure 8.**
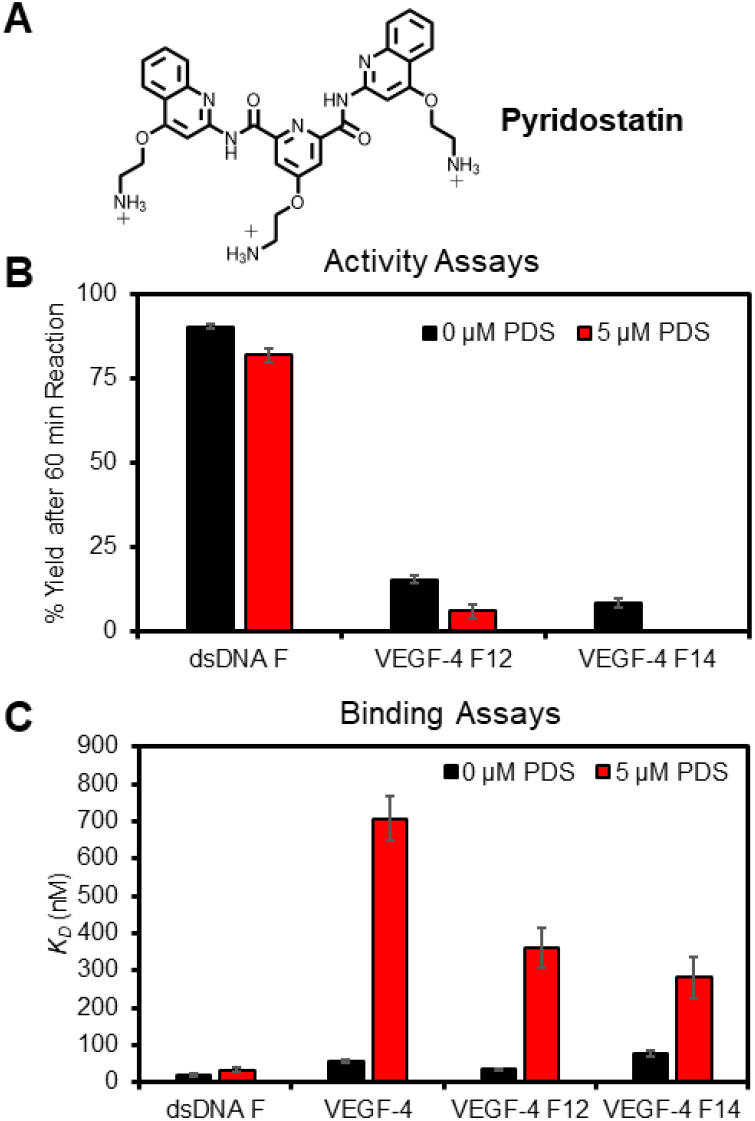
The G4 ligand PDS impacts APE1 endonuclease activity and D210A-APE1 binding of *VEGF* G4 DNA strands. (A) The structure of PDS. (B) The cleavage yields were measured after a 60 min reaction, and (C) the binding constants for the *VEGF* sequences studied. n.r. = no reaction

### 2.7 The cosolvent PEG enables APE1 to remodel F-containing dsDNA to G4 DNA

The Sugimoto laboratory has reported the properties of DNA and its interactions with proteins can change with the addition of the cosolvent polyethylene glycol 200 (PEG-200) to the buffer; they propose PEG-200 functions to mimic the molecular crowding that occurs as a result of the high concentration of proteins in the cellular milieu, or that could also occur via water exclusion.^60,61^ Studies were conducted following their guidelines with 20% PEG-200 added to the 50 mM Mg^2+^-containing buffer, and activity and binding assays were conducted. The APE1 activity on dsDNA F decreased by half and the *VEGF*-4 F12 or *VEGF*-4 F14 G4s were not cleaved in 20% PEG-200 (Figure 9A). The *K*_*D*_ value for D210A-APE1 binding dsDNA F increased twofold when 20% PEG-200 was present in a buffer (Figure 9B). In contrast, the *K*_*D*_ values found for the F-containing G4s and D210A-APE1 interactions remained nearly the same in 0% vs. 20% PEG-200 (Figure 9B). A similar result with PEG-200 was found for the five-track F-containing *VEGF* G4 folds (Figure S6). Lastly, the *K*_*D*_ value for the D210A-APE1 native *VEGF*-4 interaction was slightly diminished in the presence of 20% PEG-200 via a twofold increase when the cosolvent was present (Figure 8B). These studies reveal that the PEG cosolvent impacted the APE1 interaction with the *VEGF* sequences.

**Figure 9.**
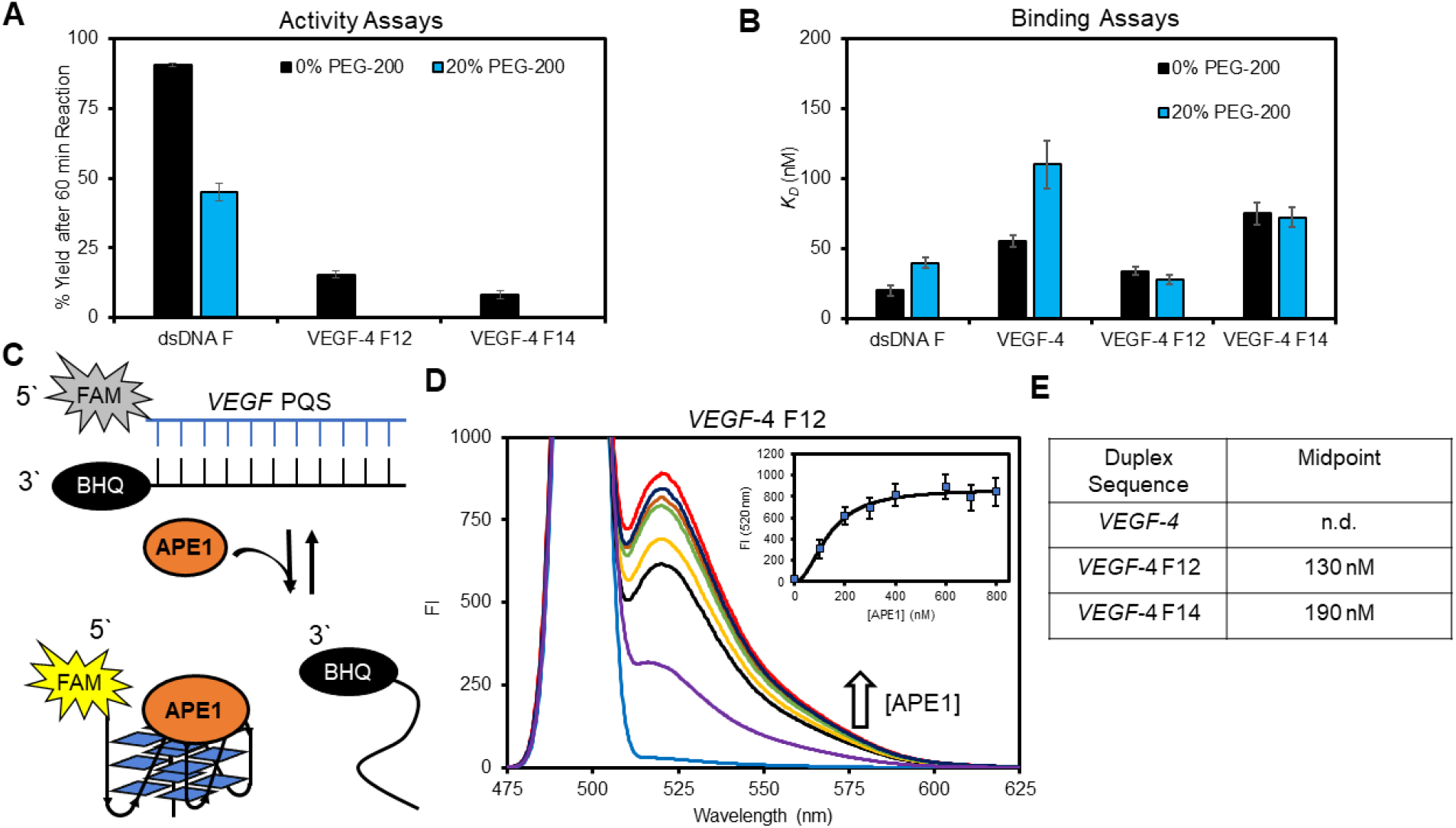
The presence of 20% PEG-200 alters the APE1 *VEGF* G4 interactions to favorably allow the protein to remodel the DNA from dsDNA to G4 DNA. (A) The APE1 activity yields determined 60 min post-incubation with and without PEG-200 present. (B) The binding of D210A-APE1 to the *VEGF* substrates with and without PEG-200 present. (C) Schematic of the FRET assay to monitor D210A-APE1 remodeling of dsDNA to G4 DNA. (D) Example FAM fluorescence spectra recorded as D210A-APE1 was titrated into a solution of *VEGF*-4 F12 dsDNA; the inset proves the fluorescence intensity at 520 nm that was fit to determine the binding midpoint. (E) The binding midpoints were obtained from the FRET assay for the *VEGF* sequences studied.

In the final study, we were curious whether APE1 could drive remodeling of dsDNA to a G4 fold because the binding was similar or slightly more favorable with the G4 in some of the binding studies conducted (Figure 9B). Remodeling of the structure was interrogated by a FRET assay in which the PQS strand was 5′ FAM-labeled and the complementary strand was labeled with a black-hole quencher (BHQ) on the 3′ side. Therefore, in the dsDNA state, FAM fluorescence is minimal upon excitation and will significantly increase upon conversion of the dsDNA to G4 DNA that releases the BHQ-containing complement (Figure 9C). The first studies were conducted without PEG-200 and no change in the FAM fluorescence was observed. Based on prior reports that found the presence of 20% PEG-200 decreased the *T*_*m*_ values for dsDNA while increasing the stability of G4 folds with F sites,^62,63^ we reasoned the presence of PEG-200 could provide a change in the thermodynamics of the system to favor the G4 fold when APE1 was present. When the FRET assays were conducted with the *VEGF*-4 F12 sequence in the presence of PEG-200, an APE1 dose-dependent increase in the FAM fluorescence was observed (Figure 9C). Fitting of the hyperbolic curve found a midpoint for the *VEGF*-4 F12 dsDNA→G4 remodeling occurred at 130 nM APE1 (Figures 9D inset and 9E). The study was repeated with the *VEGF*-4 F14 sequence to find the dsDNA→G4 remodeling occurred with a midpoint at 190 nM APE1. When the native *VEGF*-4 sequence was studied, no transition was observed that suggests the F site is needed to destabilize the dsDNA to allow the protein to refold the DNA. Thus, APE1 can function as a DNA remodeler to G4 folds. This finding aids in addressing a point in the oxidative gene regulation proposal, namely, how does the dsDNA refold to a G4 following the release of OG to yield an AP (Figure 1).

## Conclusion

Our prior work with chemically-defined plasmids studied in cellulo suggested an interplay between G oxidation in a promoter G-quadruplex sequence and the DNA repair process leading to gene induction.^1,11^ Cell-based studies guided us to propose that the unmasking of a G4 fold from the dsDNA with an embedded AP creates a hub for the endonuclease APE1 to bind but not cleave DNA. At this point, the Ref1 function of APE1 engages with activating transcription factors. Support of this proposal came from non-cleavable APE1 substrates studied in cellulo that increased the gene output from 300% to 1500%.^11^ A follow-up report supported these conclusions in genomic DNA.^64^ An unresolved question in these studies is the mechanism by which APE1 binds but does not cleave DNA in the G4 context. The present work identified many of the cellular parameters that lead to poor catalytic activity but robust binding of APE1 to the *VEGF* promoter G4. These findings may be extended to other promoter G4s, especially those that predominantly adopt parallel folds like *VEGF*.

The activity and binding assays showed that the concentrations of K^+^ and Mg^2+^ ions highly impact APE1 binding to the predominantly parallel-stranded *VEGF* G4 folds with and without an F present; moreover, in situations that resemble intracellular concentrations of these ions (140 mM K^+^ and >10 mM Mg^2+^) showed that APE1 binding the F-containing G4 is competitive or outcompetes binding with the same sequence in dsDNA (Figures 4 and 5). Additionally, when the concentration of K^+^ is 140 mM, APE1 repairs F in dsDNA but fails to process F in G4 contexts (Figure 5A). This salt influence on APE1 could be the cellular property that defines when repair of AP in dsDNA versus APE1 binding an AP in G4 DNA for gene regulation occurs. The Mg^2+^ concentration was key to significantly increase the affinity of APE1 for the G4 folds (Figure 3). Magnesium deficiency is associated with elevated levels of poorly managed oxidative stress,^65^ which may reflect the diminished APE1-G4 interaction to regulate genes responsible for responding to stress.^33,66^ Tight binding of APE1 to F-containing G4 folds requires the N-terminal domain; otherwise, the interaction has an affinity similar to F-containing ssDNA (Figures 5 and 7). Lastly, the N-terminal domain of APE1 is the region that binds the native parallel-stranded *VEGF* G4 (Figure 7). Some cancers are associated with proteasomal activity that cleaves the N-terminal tail of APE1 with dependency on the Lys acetylation status; the consequences of N-terminus removal are inappropriate localization of APE1 in the cell and altered function.^67-69^ One possible contributing factor to the cancer phenotype might be diminished APE1 binding to G4s that regulate transcription.

Experiments were conducted to probe the APE1-G4 interaction with the G4–binding agent PDS. When PDS binds the *VEGF* G4s with and without lesions, the APE1 affinity was significantly diminished (Figure 8). It is known that PDS addition to cell culture results in high levels of DNA damage that led to cell death,^70^ and blocking of APE1 binding to genomic G4s may be a contributing factor to the observed damage. Many other G4 ligands are reported for medicinal applications,^71^ and when they are bound to promoter G4s, it is possible APE1 binding will be blocked resulting in altered gene expression that may contribute to some unexpected outcomes. Thus, the design of drugs to target other features of APE1 besides G4 binding may be a good approach.^72^

A final question regarding how G4s can fold in the presence of the complementary strand favoring dsDNA was addressed in this work. For dsDNA with an F in the potential G4 context, the addition of APE1 was able to remodel the DNA from duplex to G4 (Figure 9); for this to occur, the cosolvent PEG-200 was required, which was previously proposed to function to mimic the conditions of cellular crowding^63^ or alter the water activity.^61^ As a consequence of the cellular concentration of APE1 being fairly high (10^4^-10^5^ copies/cell) with a long half-life, the concentration of the protein is sufficiently high to reach the concentration of the midpoint in the dsDNA→G4 refolding (∼150-200 nM, Figure 9).^48,73,74^ This observation aids in providing molecular understanding by which APE1 remodels F-containing promoter PQSs to G4 folds for gene induction in the cell.

Other questions remain. The binding of APE1 to native, non-lesion-containing G4s had been suggested for the human telomere sequence,^30^ but the observation that the protein binds promoter G4s is quite intriguing, especially because it is a high-affinity interaction that is similar in affinity to traditional transcription factors.^75^ Future structural studies are needed to address how the N-terminal domain of APE1 binds parallel-stranded G4s like *VEGF*. Additionally, can cellular APE1 trap G4 folds in the genome without the presence of AP sites, and what are the cellular consequences? The findings herein regarding APE1 interacting with promoter G4s for gene regulation will impact future biological and medicinal chemistry applications that target G4s with designed therapeutic drugs.

## Supporting Information

Supporting Information is available

Complete experimental methods, CD spectra, APE1 endonuclease controls studies, and endonuclease and binding data with the five G-track *VEGF* G4s

## Acknowledgments

The research was supported by the National Cancer Institute via grant no. R01 CA093099 and by the National Institute of General Medical Sciences grant no. R01 GM129267. The authors are grateful to Dr. Carla Omaga who conducted initial studies on this topic that inspired the present work.

