## Supplementary Data for "Binding of AP endonuclease-1 to G-quadruplex DNA depends on the N-terminal domain, Mg^2+^ and ionic strength"

**Item** **Page**

**Materials and Methods** S2

**Figure S1.** Table of APE1 activity conditions and the product yields obtained. S7

**Figure S2**. Example EMSA and SPR data obtained for analysis of the APE1 binding the G4s. S8

**Figure S3**. CD spectra for 5′-FAM labeled G4s compared to non-labeled folds. S10

**Figure S4**. Denaturing PAGE analysis post reaction to show demonstrate catalytic inactivity for the D210A mutants of APE1. S11

**Figure S5**. Binding plots for the F-containing *VEGF*-5 G4s. S12

**References** S13

***Materials and Methods***

***Oligomer preparation***

All DNA oligomers were synthesized and deprotected by the DNA/Peptide core facility at the University of Utah following standard protocols. The site-specific introduction of the abasic site analog tetrahydrofuran (F) was achieved using a commercially available phosphoramidite. The crude oligomers were purified using an anion-exchange HPLC column running a mobile phase system consisting of A (1 M lithium chloride, 20 mM lithium acetate at pH 7 in 1:9 MeCN:dd$H_2O$ water) and B (1:9 MeCN:dd$H_2O$). The method was initiated at 20% B and increased to 100% B via a linear gradient over 30 min with a flow rate of 1 mL/min while monitoring the absorbance at 260 nm. The purified samples were dialyzed in dd$H_2O$ for 36 h to remove the purification salts, and then lyophilized to dryness and resuspended in dd$H_2O$. The concentrations of the samples were determined by measuring the absorbance at 260 nm and using the nearest neighbor approximation to obtain the extinction coefficient for calculating the concentrations of the purified stock DNA oligomer samples. The extinction coefficients for the F-containing oligomers were estimated by omitting a nucleotide for the site. The DNA strands were annealed in the specified concentrations and buffers stated below, by heating them to 90 ˚C for 5 min in a water bath followed by gradual cooling of the water bath to room temperature.

***Circular Dichroism Analysis***

For the CD analysis, the G4 samples were annealed at a 10 μM concentration in one of three different buffer systems [(1) 20 mM Tris pH 7.4 at 22 °C with 50 mM LiOAc; (2) 20 mM Tris pH 7.4 at 22 °C with 50 mM KOAc; or (3) 20 mM Tris pH 7.4 at 22 °C 50 mM KOAc and 10 mM $Mg(OAc)_2$]. The samples were placed in 0.2-cm quartz cuvettes for CD analysis at 22 °C. The CD spectra were recorded from 220-320 nm. The spectra for the DNA strands had the background-subtracted and then the ellipticity values obtained were converted to molar ellipticity values for visualization by plotting [θ] on the y-axis and wavelength (nm) on the x-axis.

***Thermal Melting Analysis***

For thermal melting ($T_m$) experiments, oligonucleotides were first annealed at a 5 μM DNA concentration and then UV spectra were recorded in two different 50 mM $K^+$ buffer systems [(1) 20 mM $KP_i$ buffer at pH 7.4 at 22 °C with 30 mM KOAc; or (2) 20 mM $KP_i$ buffer at pH 7.4 at 22 °C 30 mM KOAc and 10 mM $Mg(OAc)_2$]. The samples were placed in a quartz Tm analysis cuvette that was placed in a temperature-regulated UV-vis spectrometer followed by thermal equilibration at 20 °C before the commencement of the experiment. The thermally induced denaturation of the G4s was monitored at 295 nm by heating the sample from 20 to 100 °C at a ramp rate of 0.5 °C/min followed by a 60-sec equilibration and then measuring the absorbance value at 295 nm. The absorbance data collected were background subtracted and then $T_m$ values were determined using Shimadzu's $T_m$-analysis software.

***Expression and purification of recombinant APE1***

Human wild-type APE1 was expressed from the pET28HIS-hAPE1 plasmid purchased from Addgene (#70757).[1] The D210A-APE1 mutant that is catalytically inactive was constructed using the Q5 Site-Directed Mutagenesis Kit (NEB). The D210A-APE1 truncate mutant plasmids for Δ33-D210A-APE1 and Δ61-D210A-APE1 were prepared using the Q5 Site-Directed Mutagenesis Kit (NEB). All mutated plasmids were verified by Sanger sequencing.

The APE1 proteins were expressed in BL21(DE3) competent *E. coli* cells (NEB), grown at 37 °C until induced at OD = 0.6 with 100 μM IPTG, and then grown overnight at 37 °C. After harvesting by centrifugation, the cells were lysed by sonication in 20 mM sodium phosphate, 300 mM sodium chloride (pH 7.4) with 1 mM PMSF, and 5 mM BME. The lysate was pelleted at 18,000 × g for 30 min. The resulting supernatant was passed through HisPur™ Ni-NTA resin (Thermo Fisher Scientific) equilibrated with the lysate buffer. The proteins were eluted from the Ni-NTA column with a linear gradient of imidazole from 10 mM up to 250 mM. The proteins eluted at high imidazole concentration were buffer exchanged into 10 mM sodium phosphate, 50 mM NaCl, 1 mM DTT, 50% glycerol (pH 7.4 at 25 ˚C). The resulting buffer exchanged proteins were stored at −20 °C. The final concentrations were determined by a NanoDrop One UV-vis spectrophotometer.

### *DNA substrate preparation for APE1 activity assays*

The F-containing VEGF DNA strands were $^{32}P$ radiolabeled by allowing the strands to react with $^{32}P$-ATP in the presence of T4-polynucleotide kinase (NEB) following the manufacturer's protocol. The radiolabeled DNA strands were purified using PD Spin-Trap™ G-25 column (Cytiva) following the manufacturer's protocol. Reaction solutions were made with 4 nM $^{32}P$-labeled DNA mixed with 6 nM non-labeled DNA. The G4 folds from the mixed sample were prepared in the various buffers outlined in Table 1 below. The dsDNAs were prepared identically to the G4 samples with the addition of 1.5-fold excess of the complementary strand before annealing.

**Table 1**. Buffer conditions studied

| [Tris] (mM) | [KOAc] (mM) | [LiOAc] (mM) | [$Mg(OAc)_2$] (mM) | %PEG-200 |
|---|---|---|---|---|
| 20 | 50 | | 10 | |
| 20 | 140 | | 10 | |
| 20 | | 50 | 10 | |
| 20 | 50 | | 0 | |
| 20 | 50 | | 1 | |
| 20 | 50 | | 10 | 20 |

### *APE1 activity assays*

Using the prepared samples from the previous step, the reactions were conducted in 10-µL reactions by pre-incubating the 10 nM DNA samples at 37 °C for 30 min. After the incubation, APE1 was added to a 3 nM final concentration, and the reactions were incubated at 37 ˚C for up to 60 min. Quenching the samples at the desired time point was achieved by adding EDTA to 10 mM concentration and an equal volume of stop solution (95% formamide, 5% glycerol with 0.1% each of xylene cyanol and bromophenol blue) followed by heat denaturing the samples at 65 °C for 20 min. Assay mixtures without enzymes were used as negative controls. The samples were then analyzed by placing them in a 20% 7 M urea-denaturing PAGE followed by electrophoresing them at 75 W for 2.5 h. The gels were exposed on a storage phosphorimager screen for 18 h and the bands were visualized by autoradiography (TyphoonTM 9400 Variable Mode Imager (GE Amersham Biosciences)) followed by band quantification using ImageQuant™ image analysis software. The percent yields were calculated by dividing the band intensity of the product band by the sum of the product and reactant bands and then multiplied by 100. The standard deviations obtained from triplicate trials of each reaction provided the error bars for the data reported.

***DNA substrate preparation for fluorescence anisotropy binding assays***

The fluorescence anisotropy assays were conducted with VEGF DNA strands labeled on the 5′ end with 6-carboxyfluorescein phosphoramidite (FAM). The 5′-labeled DNA strands were annealed at 100 nM concentration in the buffers outlined in Table 1 following the procedure previously described.

***Fluorescence anisotropy binding assays***

The fluorescence anisotropy experiments were conducted by titrating the prepared 5′-FAM-labeled DNA samples with the appropriate APE1 protein from 0-5000 nM followed by incubating the mixture at 22 °C for 30 min before analysis. The fluorescence anisotropy measurements were carried out on a BioTek Synergy2 Multi-Mode Microplate Reader at excitation and emission wavelengths 485 and 520 nm, respectively. The anisotropy values (r) were calculated with Eq. 1 in which $I_{par}$ is the parallel emission intensity and $I_{per}$ is the perpendicular emission intensity.

$$r = \frac{I_{par} - I_{per}}{I_{par} + 2I_{per}} \qquad \text{Eq. 1}$$

The r values obtained were plotted against log[APE1] to produce sigmoidal curves that were fit to the following Hill equation where the bottom is the lower plateau value of r and the top is the higher plateau value of r defining the sigmoidal curve. The dissociation-binding constant is $K_D$ and *n* is the Hill coefficient (Eq. 2).[2]

$$\mathrm{r} = bottom + \frac{(top - botto\quad)}{1 + 10^\wedge(logKd - \log[APE1])^{n}} \qquad \text{Eq. 2}$$

The $K_D$ and n values were determined from the equation obtained that best fit the data points based on linear least-squares analysis. The error bars in the values reported are the standard deviations of the values obtained from triplicate trials.

***FRET assay***

The fluorophore-quencher duplex DNA samples were prepared using the same conditions as described for the APE1 cleavage assays, with the exception that the DNA concentrations were 100 nM. In analyses that incorporated PEG-200, the cosolvent was added to a 20% concentration and all other conditions remained the same. Samples of DNA at 200 µL were incubated with a concentration series of APE1 from 0-800 nM and incubated at 37 °C for 30 min before fluorescence analysis. The fluorescence spectrometer scanned the emission spectrum from 450-625 nm with an excitation wavelength of 495 nm. Plots of the emission intensity at 520 nm vs. [APE1] were constructed to follow the duplex to G4 equilibrium shift.

**Figure S1.** Table of APE1 activity conditions and the product yields obtained.

| [VEGF-5 F14] (nM) | [APE1] (nM) | %Yield |
|---|---|---|
| 10 | 3 | ~20 |
| 10 | 10 | ~20 |
| 10 | 30 | ~20 |

The other *VEGF* G4s were studied under the variable conditions to find no change in the yields upon changing the reaction conditions.

**Figure S2**. Example EMSA and SPR data were obtained for analysis of the APE1 binding the G4s.

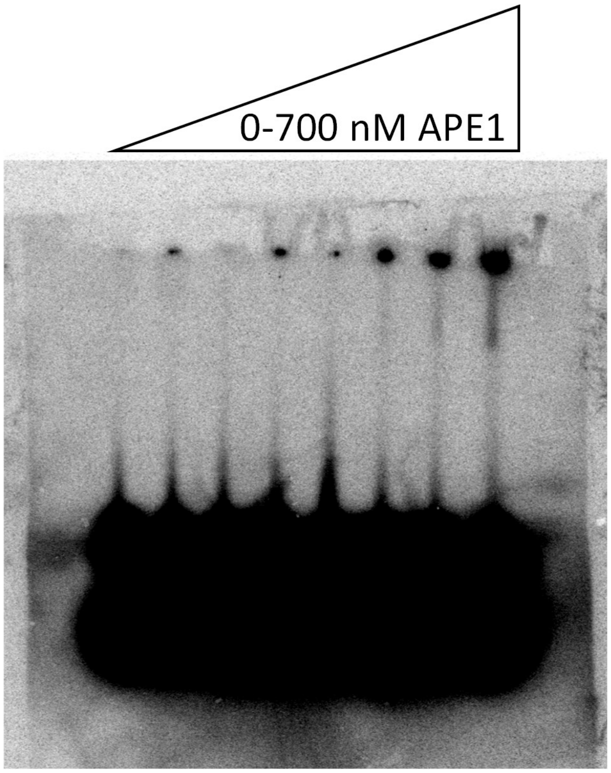


Example EMSA for analyzing D210A-APE1 binding *VEGF*-4 F12. The material would not migrate out of the wells to give a bound band vs. unbound band for quantification. In our hands, this approach to monitor the protein G4 interaction could not be used for quantitative analysis.

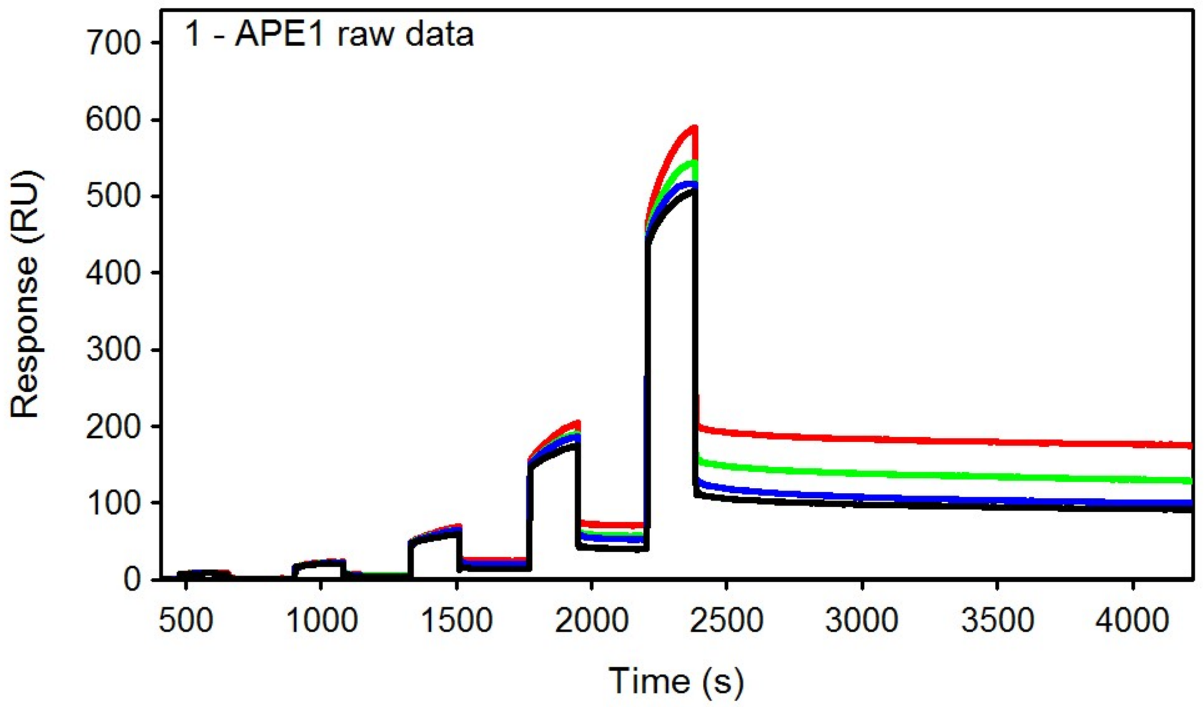


SPR sensorgrams for a three-fold titration series up to 300 nM APE1 illustrating non-specific binding to the sensor chip because the signal does not return to the base line value after each APE1 addition to the flow cell. The data were collected in a buffered solution comprised of 50 mM Tris (pH 7.4), 50 mM KOAc, and 1 mM DTT at 25 °C with *VEGF*-4 F12 surface bound and a three-fold titration series of APE1 up to 300 nM.

**Figure S3**. CD spectra for 5′-FAM labeled G4s compared to non-labeled folds.

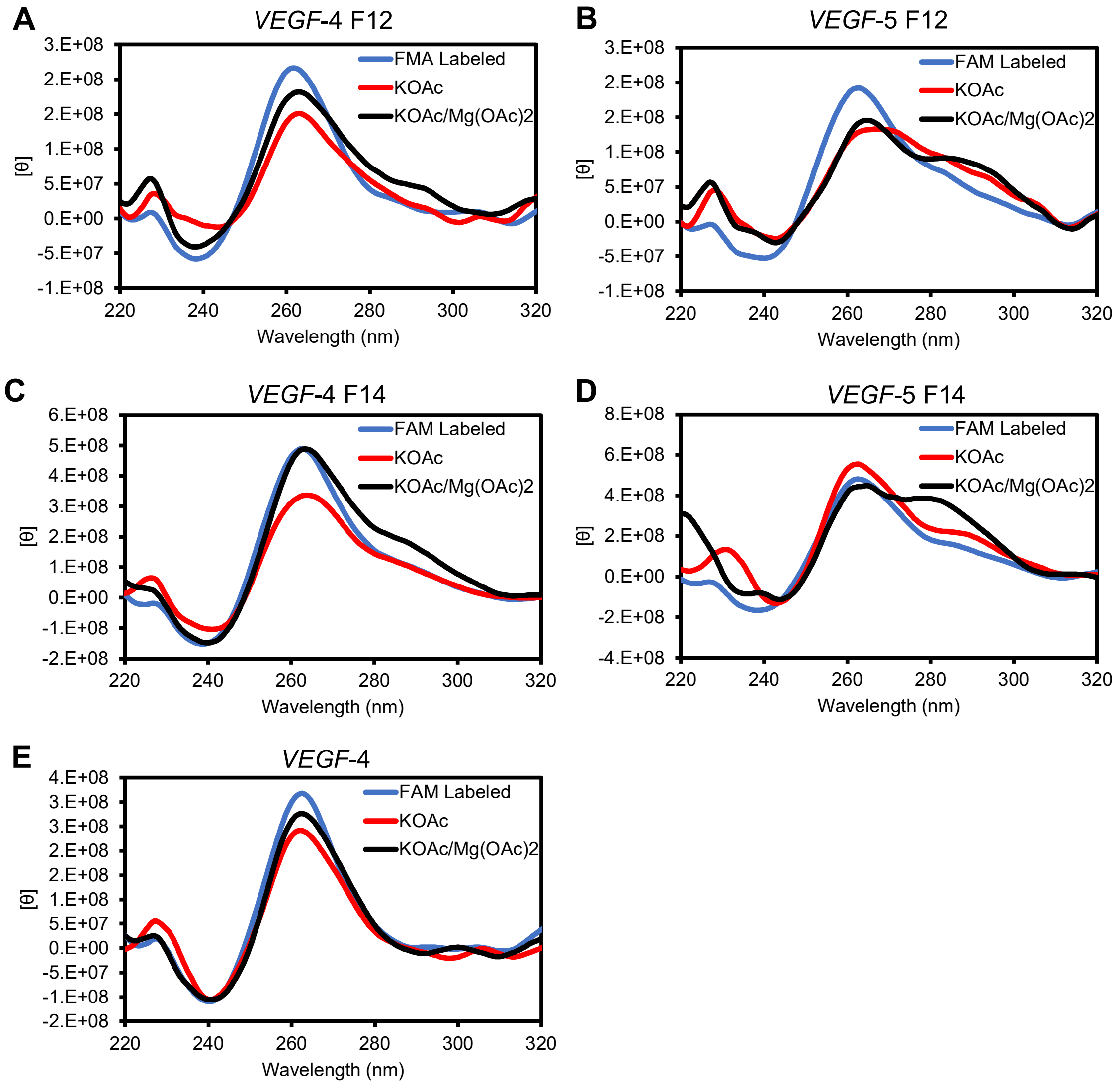


The FAM-labeled VEGF G4s were annealed in 20 mM Tris pH 7.4 at 22 °C with 50 mM KOAc present before recording the CD spectra (blue line). Spectra for each sequence are as follows (A) *VEGF*-4 F12, (B) *VEGF*-5 F12, (C) *VEGF*-4 F14, (D) *VEGF*-5 F14, and (E) *VEGF*-4.

**Figure S4**. Denaturing PAGE analysis post reaction to show demonstrate catalytic inactivity for the D210A mutants of APE1.

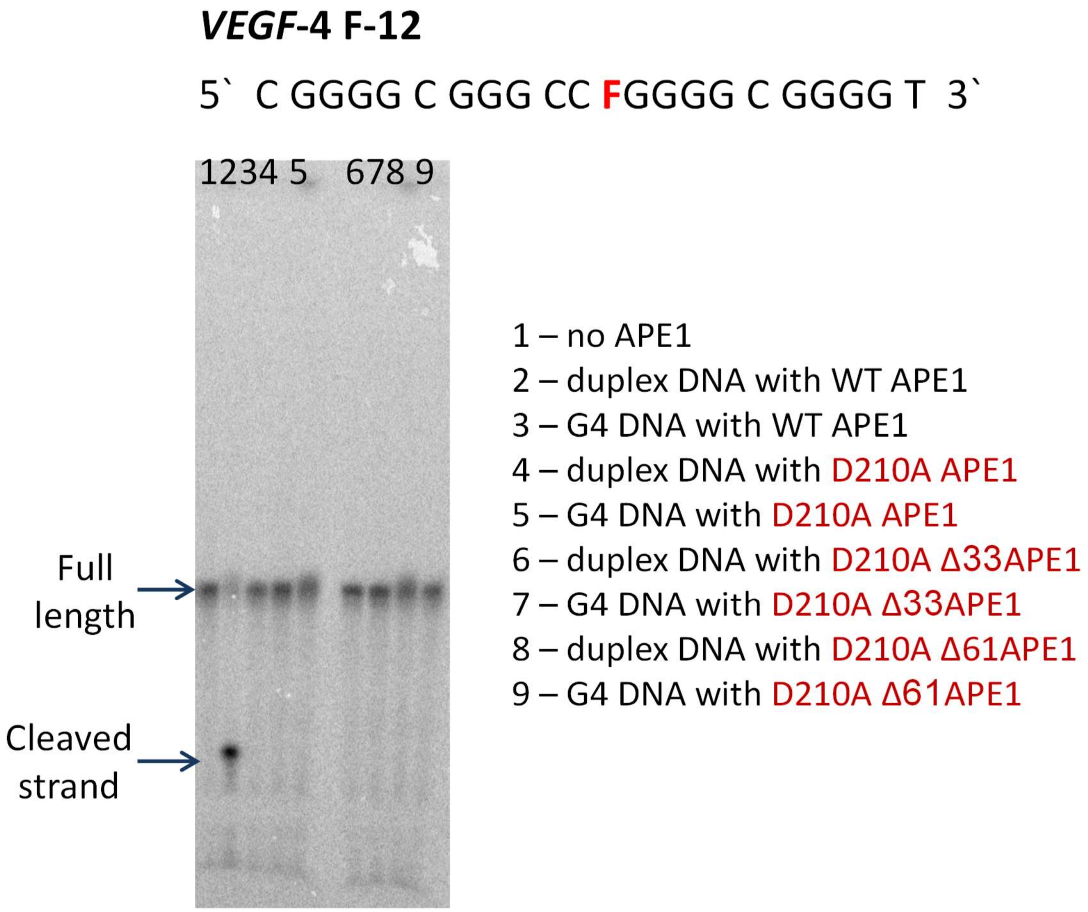

**Figure S5**. Binding plots for the F-containing *VEGF*-5 G4s.

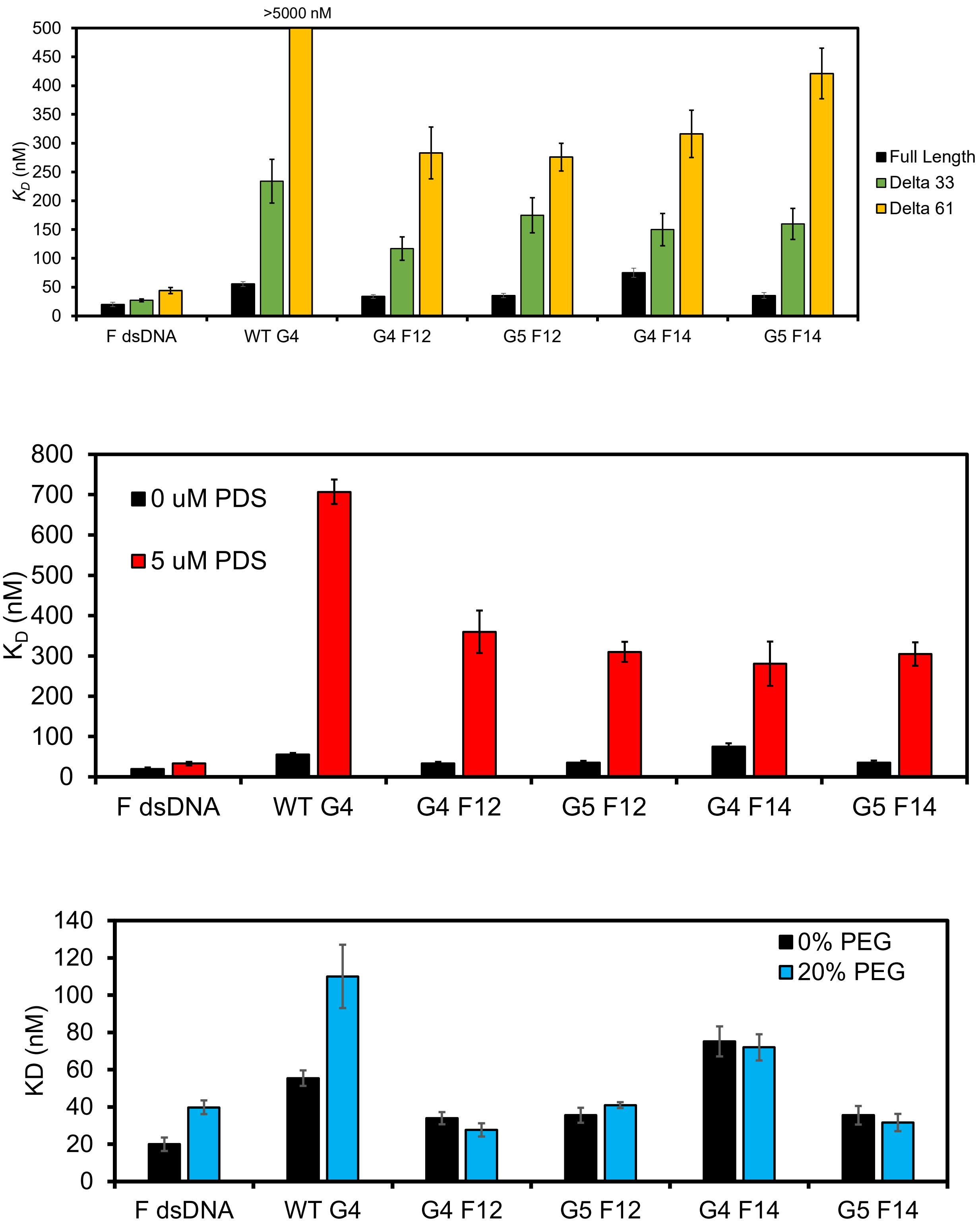
